# Detecting epistatic interactions in genomic data using Random Forests

**DOI:** 10.1101/2022.04.26.488110

**Authors:** Hawlader A. Al-Mamun, Rob Dunne, Ross L. Tellam, Klara Verbyla

## Abstract

Epistatic interactions can play an important role in the genetic mechanisms that control phenotypic variation. However, identifying these interactions in high dimensional genomic data can be very challenging due to the large computational burden induced by the high volume of combinatorial tests that have to be performed to explore the entire search space. Random Forests Decision Trees are widely used in a variety of disciplines and are often said to detect interactions. However, Random Forests models do not explicitly detect variable interactions. Most Random Forests based methods that claim to detect interactions rely on different forms of variable importance measures that suffer when the interacting variables have very small or no marginal effects. The proposed Random Forests based method detects interactions using a two-stage approach and is computationally efficient. The approach is demonstrated and validated through its application on several simulated datasets representing different data structures with respect to genomic data and trait heritabilities. The method is also applied to two high dimensional genomics data sets to validate the approach. In both cases, the method results were used to identify several genes closely positioned to the interacting markers that showed strong biological potential for contributing to the genetic control for the respective traits tested.

## 1 Introduction

Many phenotypes and disease traits in human, animals and plants are complex and involve many genes and their interactions. Quantitative traits such as human height or yeast growth are influenced by many genetic variants each typically of small effect size. Numerous genetic variants have been reported in genome-wide association studies (GWAS) and quantitative trait loci (QTL) studies that are associated with various quantitative traits in different species. However, for each trait, the reported variants cumulatively can explain only a portion of the total heritability of the trait. This phenomenon of unexplained heritability is termed “missing heritability” (Maher, 2008), and is partly due to not considering the genetic variant interactions (epistasis) between the genetic markers (Zuk et al., 2012).

Epistasis occurs when there is any non-additive interaction relationship between two or more genes in their combined effects on a phenotype. It can be potentially identified when the effects of two or more genetic markers associated with a trait differ from the sum of the individual marker effects. Understanding epistasis is important because it can help explain the functional mechanisms of the genes that together contribute to disease or trait expression and enable more targeted and nuanced interventions to be developed. For example, being able to estimate the effect of non-additive interactions could potentially be exploited to improve the genetic predictions (Ansarifar et al., 2020) for economically important traits such as wheat yield or carcass weight of cattle.

The most direct approach to detect epistatic interactions is to use linear models (LM) to evaluate all pair-wise loci combinations to test for non-additive interactions associated with a complex trait (Zhang et al., 2008). This type of exhaustive search is only feasible when the number of candidate markers is relatively small or the computational power available is extreme. However, as the number of markers in a dataset grows, this approach suffers from the computational burden induced by the combinatorial number of tests that have to be performed. For example, The Welcome Trust Case Control studies (Burton et al., 2007) had 17,000 individuals (2,000 cases for each of seven diseases and 3,000 combined controls) genotyped with the GeneChip 500K Mapping Array Set (Affymetrix chip), which interrogated 500,568 SNPs. Thus, more than 1.25 × 1013 pairwise tests would need to be performed, which is not computationally feasible using standard equipment. Moreover, recent technological advancements and reduced costs have led to experimental designs utilising more than 1 million variants and markers for hundreds of individuals, which massively confound the ability to identify interacting pairs of SNPs (Ha et al., 2014; Gholami et al., 2014; Auton et al., 2015; University of Utah, 2021). Conse-quently, two step-based methods have been proposed to detect epistatic interactions from genomic data, where in the first step markers were pre-selected or weighted and then in the second step interaction testing was performed only on the reduced set of markers. The preselection can be performed based on statistical tests (e.g., GWAS (Pecanka et al., 2017)) or machine learning methods with feature selection (Jiang et al., 2009; Meng et al., 2007).

Random Forests algorithms are an ensemble learning method first proposed by Leo Breiman (Breiman, 2001). Ensemble learning combines multiple learning algorithms/models to improve prediction or classification performance beyond what could be obtained from the constituent models. In Random Forests, *m* random samples (with replacement) are selected by bootstrap sampling. Then multiple decision trees are created based on the bootstrap samples. To break down the correlation between features, only a random subset of features is considered at any node of the tree building process. For classification problems, the class of the validation data is predicted based on majority voting and for the regression problem, the model outputs the mean of the individual tree’s prediction. There are two main parameters in the implementation of the Random Forests algorithm, namely the number of decision trees (*num*.*trees*) and the number of randomly selected features (*mtry*) from which the best feature for splitting is selected at any given node.

Random Forests models are widely used (Lundberg et al., 2019) in a variety of disciplines. The attraction of the model is based on several attributes:

- a high degree of accuracy;
- a resistance to overfitting, including in the presence of large

numbers of redundant variables;

- the provision of an out-of-bag error measure;
- the provision of a variable ranking at little extra computational cost.

It is clear that Random Forests (RF) can perform well with correlated and interacting variables, at least in the sense that the fitted model performs well (Ziegler et al., 2007). However, most Random Forests based approaches are either based on single variable importance (Jiang et al., 2009; Yoshida and Koike, 2011) or based on pairwise variable importance (Ziegler et al., 2007). The Random Forests-based approaches for identifying interactions based on variable importance measurements cannot distinguish whether a discovered interaction is a true interaction or simply two variables with strong marginal effects. If the interacting variables have small marginal effects, they will not be highly ranked on the variable importance list such that detecting interactions based on variable importance metrics can be problematic. Variable importance measurements are also heavily dependent on extensive permutation tests. Schmalohr et al. (2018) notes that there is a lack of accurate methods for the detection of significant interactions between genomic loci but that several studies have shown that Random Forests can outperform other QTL mapping methods when identifying QTL or marker trait associations. They also provided benchmarks on simulated and real yeast cross data indicating that the Random Forests-based methods outperform other commonly used approaches for detecting epistasis.

To overcome the challenge of a large number of pairwise interaction tests, we proposed an effective two-step process based on our observations from using a Random Forests algorithm. Using simulation data, it was observed that if two markers are interacting, they appear as parent-child nodes in a decision tree of a Random Forests more frequently than if they are not interacting. This new method and the validation results obtained using simulated data are presented in this manuscript. In addition, the results from the use of the method using two real datasets are discussed.

## 2 Methods

The proposed method is described in Figure 1. First, a Random Forests model is created using all available genomic data. For both the simulated and real data examples, the R package *ranger* (Wright and Ziegler, 2017) was used with the parameters *num*.*tree* = 1000 and *mtry* = #SNPs/3.

**Figure 1.**
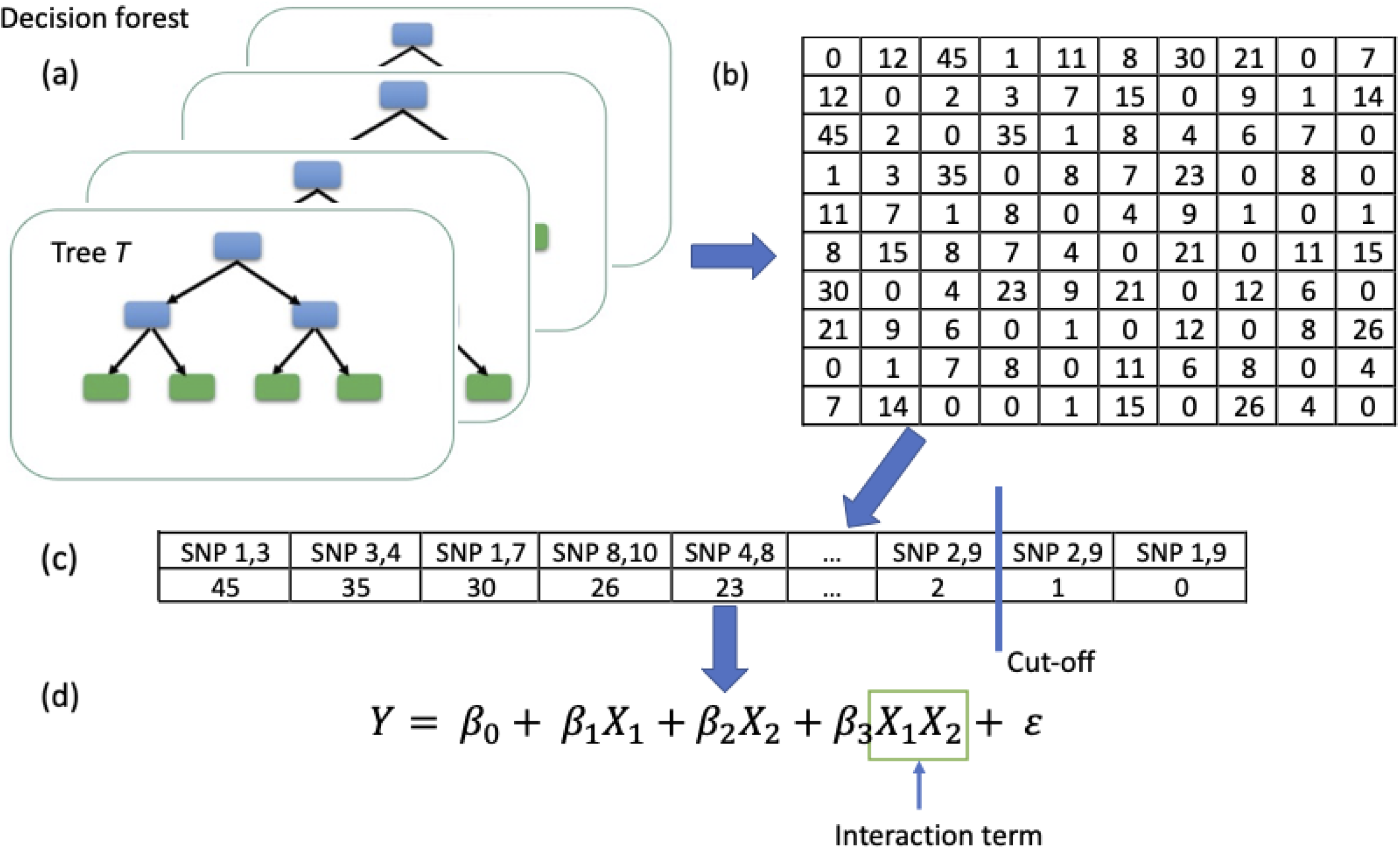
The stages of the proposed method. (a) A Random Forests model is trained with all genomic markers. (b) A parent-child node count matrix is generated. (c) Marker pairs are ranked based on the number of appearances as parent-child pairs. (d) Using linear model, interaction tests are performed only for the selected marker pairs.

Once the Random Forests model is built, the number of times each pair of markers was found as parent-child nodes in all the decision trees were recorded. Then the marker pairs were ranked based on the count and only the pairs having count more than 2 were retained. A threshold of at least two (*t* = 2) appearances as parent-child nodes in the decision trees of the Random Forests was used because the use of one (*t* = 1) resulted in a substantially higher number of SNP pairs, often random occurrences, that did not improve the identification of interactions.

Finally, using linear models the significance of the interactions are tested. For a linear model, an interaction occurs when an independent variable has a different effect on the outcome depending on the values of another independent variable. The variables in the model are divided between main and interaction effects. A multiplicative epistasis model was used, where the interaction between two markers, *i* and *j*, are modelled as the product of the two genotype values, using the following linear regression model:

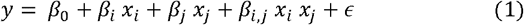

where *y* is the quantitative measure of phenotype, *x*_*i*_ and *x*_*j*_ are the genotype values {0, 1, 2} for SNP *i* and *j* respectively, *β*0 is the mean phenotype, *β*i and *β*j are the marginal effects for SNP *i* and *j, β*_i,j_ is the interaction effect between SNP *i* and *j*, and ε is the residual with N(0,1). For case-control disease data, the interaction was modelled through a logit function as follows:

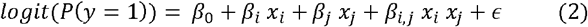

where ‘1’ is the disease state and ‘0’ is the control state. An interaction is considered identified when the *p*-value of *β*i,j is significant.

### 2.1 Simulation data

Three different interaction scenarios were considered to access the performance of the proposed method in detecting pairwise interactions.

A simulated dataset was generated using publicly available genotype data containing of 3,534 animals from a single PIC nucleus pig line genotyped using the Illumina ProcineSNP60 chip (Cleveland et al., 2012). The original dataset contained 52,843 single nucleotide polymorphisms (SNP) but in this study, all non-informative SNPs were removed leaving 48,015 markers. The simulated data was created by randomly selecting 5,000; 10,000 and 48,000 markers from the dataset.

Using these three datasets, simulated phenotypes were then created having different heritabilities (*h*2 = 0.25, 0.33, 0.50, 0.66, 0.75 and 1.00). Data was generated using three different models, a “Base model” where none of the markers nor their interactions have any effect on the phenotype; a “No Interaction model” where markers have marginal effects but there is no additional interaction effect, and in the “Interaction model” where three pairs of markers have no marginal effects but have interaction effects. The scenarios can be summarised as:

- **Base model** phenotype = sum of ten QTL effects + noise (depending on heritability);
- **No Interaction model** phenotype = sum of ten QTL effects + sum of marginal effect of epistatic SNPs (no interactions) + noise (depending on heritability);
- **Interaction model** phenotype = sum of ten QTL effects + sum of interaction effects for the three pairs (no marginal effects) + noise (depending on heritability).

### 2.2 Real dataset – Wellcome Trust Case Control Consortium data for rheumatoid arthritis

The first real dataset used was obtained from the Wellcome Trust Case Control Consortium (WTCCC) (Burton et al., 2007). The original dataset contains 14,000 cases for seven complex human diseases and 3,000 shared controls. The control individuals were from two different sources: 1958 British Birth Cohort (58C) and the UK Blood Service Control Group (UKBS). Each control group has 1,500 individuals. Samples of DNA from all 17,000 individuals were genotyped with the GeneChip 500K Mapping Array Set (Affymetrix chip). The current analysis only used the data for rheumatoid arthritis (RA) that consisted of 4,798 individuals with 1,860 cases and 2,938 controls. SNP variants with more than 10% of calls missing and MAFs of less than 0.05 were removed. SNP markers that failed the Hardy-Weinberg test (p ≤ 0.05) were also removed. As linkage disequilibrium (LD) could affect how the Random Forests (RF) algorithm selects a SNP at a particular node (for more detail on how LD can affect RF, please see (Meng et al., 2009)), the dataset was also filtered based on LD using PLINK 1.9 (Purcell et al., 2007). Filtering removed one SNP from a pair of SNPs within a 200 kb window if pairwise LD exceeded the threshold *r*2 >= 0.3. The final dataset contained 94,484 SNP markers and 4,798 individuals.

### 2.3 Real dataset – Mouse data

The second real data set used was published by (Solberg et al., 2006; Valdar et al., 2006). A curated version of this dataset was downloaded (Additional file 1 in Martini et al. (2017)) and contained 9,265 SNPs for each of 1,298 individuals. Pre-corrected residuals of thirteen phenotypes were available as phenotypes. In the current analysis, the phenotype “weight at six weeks” (W6W) was used.

## 3 Results and Discussion

### 3.1 Simulation data

The method’s performance was assessed by determining the number of interactions that were correctly identified and through evaluating the repeatability and false discovery rate for the approach.

The evaluation of the method’s ability to correctly identify interacting markers was assessed based on the proportion of time, the method selected the interacting marker pairs for pairwise interaction tests. Table 1 shows the importance ranking of three interacting SNP pairs based on the number of times they appeared as parent-child nodes in the Random Forests decision trees. Higher ranking, represented by a smaller number, meant a higher count. For example, a value of 1 for pair P3 means that SNP pair 3 appeared as parent-child nodes more than any other pair of markers in that Random Forests analysis.

**Table 1.** Ranking of the SNP pairs based on counts. Simulated data for 10 QTL and 3 pairs of interactions. NC = No count. P1, P2 and P3 are three interacting SNP pairs.

| Num<br>SNP | $h^2$ | Base model | | | No interaction | | | Interaction | | |
| --- | --- | --- | --- | --- | --- | --- | --- | --- | --- | --- |
|  |  | P1 | P2 | P3 | P1 | P2 | P3 | P1 | P2 | P3 |
| 5,000 | 0.33 | NC | NC | NC | 323 | 2 | 287 | 10 | 5 | 7 |
|  | 0.50 | NC | NC | NC | 151 | 1 | 3373 | 6 | 4 | 10 |
|  | 0.66 | NC | NC | NC | 730 | 3 | 699 | 8 | 4 | 9 |
|  | 1.00 | NC | NC | NC | 230 | 1 | 315 | 7 | 3 | 11 |
| 10,000 | 0.33 | NC | NC | NC | NC | 20 | NC | 132 | 9 | 3 |
|  | 0.50 | NC | NC | NC | NC | 40 | NC | 293 | 7 | 1 |
|  | 0.66 | NC | NC | NC | NC | 59 | NC | 88 | 5 | 1 |
|  | 1.00 | NC | NC | NC | 1077 | 32 | NC | 86 | 6 | 1 |
| 48,000 | 0.33 | NC | NC | NC | NC | NC | NC | 4 | 50 | 63 |
|  | 0.50 | NC | NC | NC | 204 | 412 | NC | 5 | 27 | 12 |
|  | 0.66 | NC | NC | NC | 806 | 224 | NC | 2 | 30 | 13 |
|  | 1.00 | NC | NC | NC | 663 | 118 | 909 | 13 | 45 | 9 |

As was expected in the “Base model”, three pairs of interacting SNP do not appear as parent-child nodes in any decision tree of the Random Forests. This reflected that these SNP did not have any marginal or interaction effects. In the “No interaction model”, the three interacting SNP pairs appeared as parent-child nodes in some of the scenarios. This is expected as in the “No Interaction model” all the individual SNP have their own marginal effects while not interacting. Hence, they were correctly selected at different nodes of the decision trees.

For the “Interaction model”, rankings of the three interacting SNP pairs are relatively higher than the other two models. The rankings observed demonstrated that if the SNP pairs are ranked based on the count of their appearance as parent-child nodes, it should be possible to only test pairwise interaction for a subset of marker pairs from the vast number of total possible combinations of markers.

To quantify the efficiency of the method in reducing the search space, the number of tests performed was compared to the number of markers and potential pairwise combinations. For this test, a threshold of *t* = 2 was used, meaning that if any of the marker pairs appeared as parent-child nodes in the decision trees of the Random Forests for at least twice, they were then tested for interaction significance using Equation 1. Table 2 shows the number of pair-wise interaction tests performed for the “Interaction model” alongside the number of possible pairwise combinations for specified sets of heritability and marker numbers. The final column of Table 2 shows that for all scenarios, less than one percentage of the total marker pairs were tested for interaction. In all cases, including for the largest simulated dataset with 48,000 markers, the three interacting pairs (P1, P2 and P3) were selected for pairwise interaction tests.

**Table 2.** Number of tests performed to detect interactions.

| Num SNP | Num of possible combination | $h^2$ | Num test performed | Percentage (%) |
| --- | --- | --- | --- | --- |
| 5,000 | 12,497,500 | 0.33 | 61,346 | 0.49 |
|  |  | 0.50 | 62,610 | 0.50 |
|  |  | 0.66 | 64,111 | 0.51 |
|  |  | 1.00 | 57,747 | 0.46 |
| 10,000 | 49,995,000 | 0.33 | 18226 | 0.04 |
|  |  | 0.50 | 18938 | 0.04 |
|  |  | 0.66 | 20324 | 0.04 |
|  |  | 1.00 | 21204 | 0.04 |
| 48,000 | 1.15E+09 | 0.33 | 1742 | 1.51E-04 |
|  |  | 0.50 | 2400 | 2.08E-04 |
|  |  | 0.66 | 3064 | 2.66E-04 |
|  |  | 1.00 | 5531 | 4.80E-04 |

Table 2 demonstrates that, as the number of markers increased the total number of tests performed decreased. This result is due to the fact that the number of trees (*num*.*tree* = 1000) and a threshold of *t* = 2 remained the same for all simulations. Thus, as combinatorial search space increases due to increasing numbers of markers, the number of pairs of markers that exceed the parent-child node threshold (*t* = 2) is reduced. This suggests that for smaller marker sets, the number of decision trees could be reduced to minimise computation and time demands. One strategy could be to adjust the number of decision trees dependent on the number of markers.

#### 3.1.1 Heritability

The effect of heritability on the method’s performance was explored with the pairwise counts for each of the interacting SNP pairs in each Random Forests at different heritabilities (*h*2 = 0.33, 0.50, 0.66, and 1.00) for 10,000 SNP with 10 QTL and 3 pairs of interacting SNPs recorded. The results are shown in Figure 2. At lower heritabilities, the pairwise count is small and as heritability increases, the pairwise count also increases. Although the three interacting SNP pairs have very similar interaction effects (ranging between 0.3 to 0.4), they have very different numbers of counts. There is a direct relationship between the *p*-value of the interaction term with the pairwise count. Lower *p*-values of the interaction terms have higher pairwise count (correlation coefficient = 0.84, p-values and correlations are not shown). This is particularly interesting as even very small interaction effects with high significance will not be overlooked because of higher pairwise count.

**Figure 2.**
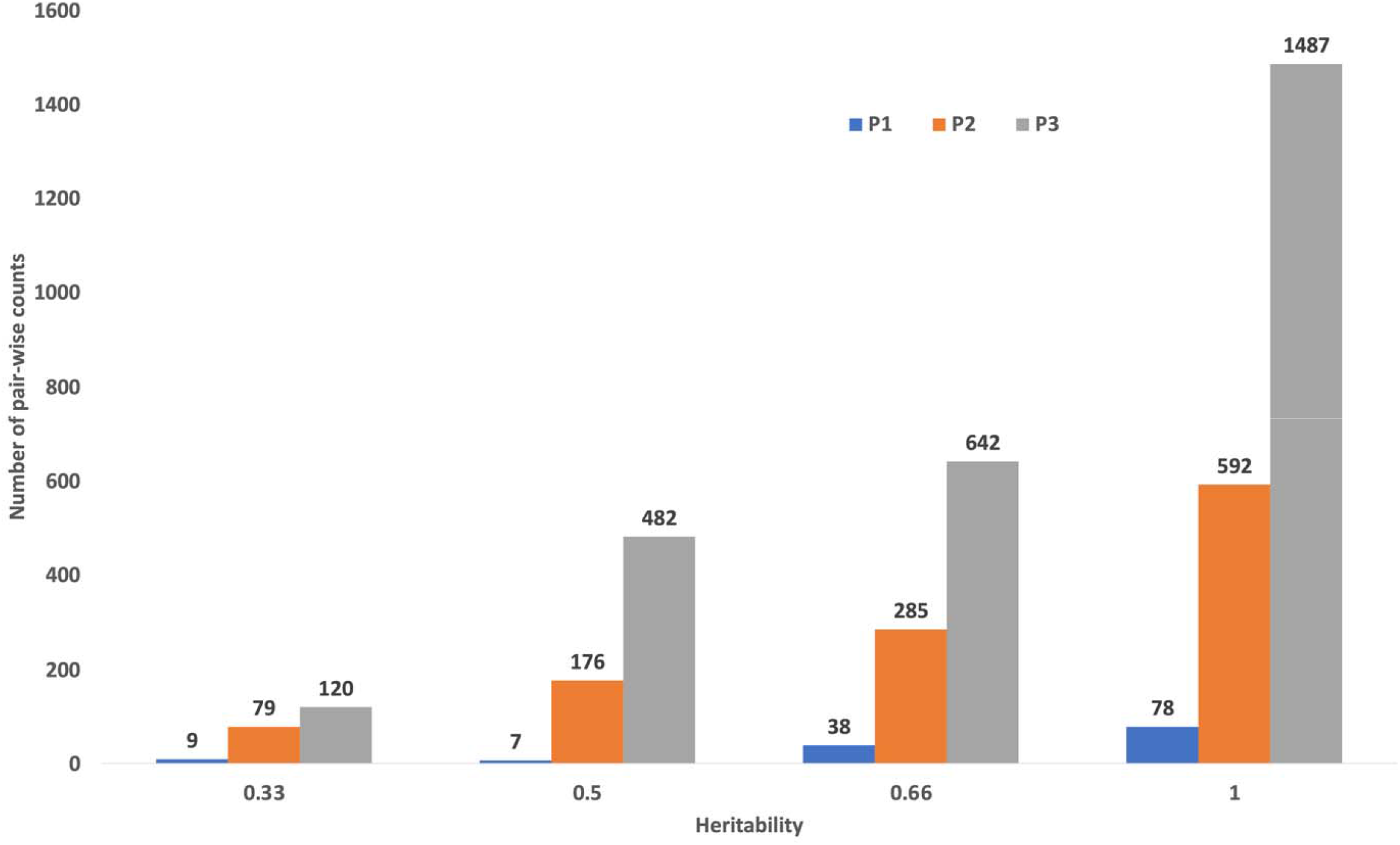
The effect of trait heritability on pairwise counts. P1, P2 and P3 are interacting SNP pair 1, 2 and 3 respectively. The Y-axis shows the number of pairwise counts for each interacting SNP pair in each Random Forests.

#### 3.1.1 Repeatability and false discovery rate

To test repeatability of the proposed method, the “Interaction model” was generated ten times at each heritability (0.25, 0.33, 0.50, 0.66, and 0.75) with the correct identification of the interacting marker pairs assessed. Table 3 shows the results for 10,000 SNPs with 10 QTLs and 3 pairs of interacting SNPs. In most of the scenarios tested, the method was able to successfully identify the interacting pairs except for pair P1 at lower heritabilities. This result appears to be because pair P1 has a lower interaction effect when compared to pairs P2 and P3. Pair P1 appeared on the candidate lists in all instances but did not pass multiple test corrections several times when the heritabilities were low (ie 0.25, 0.33).

**Table 3.** Repeatability and number of false positives for the proposed method. The data contains 10,000 with 10 QTLS and 3 pairs of interacting SNP. The process was repeated 10 times for each heritability with the number of times each pair (P1, P2 or P3) found and the mean number of false positive interactions identified indicated.

| $h^2$ | P1 | P2 | P3 | False Positive |
| --- | --- | --- | --- | --- |
| 0.75 | 10 | 10 | 10 | 0 |
| 0.66 | 10 | 10 | 10 | 0.1 |
| 0.50 | 9 | 10 | 10 | 0.3 |
| 0.33 | 7 | 10 | 10 | 0.8 |
| 0.25 | 4 | 10 | 10 | 1.9 |

To assess the false discovery rate of the method, the number of false positive interactions were counted across the 10 replicates at the different heritability levels. A false positive was declared when a pair of SNP were not in the original data and when these SNP were not correlated or in high LD with one of the three simulated pairs. This is because there were multiple interactions identified where each marker in the pair correlated highly with corresponding markers in one of the simulated interaction pairs. Only when this was not the case was a false positive declared. A Pearson correlation coefficient of less than 0.25 was used as the threshold. This threshold was chosen as using a 0.5 threshold revealed a number of pairs incorrectly identified as false positives where each marker in the pair was correlated, normally with a coefficient above 0.4, with one of the simulated three pairs. The mean false positive results for each heritability calculated using the ten replicates, are shown in Table 3. The number of false positives increased in the data where the heritability was higher. This was understandable as the variance explained by the marker data (genetic signal) increases at higher heritabilities. The number of false positives detected by the proposed method was low across all heritabilities.

### 3.2 Results for real datasets

To evaluate the method’s performance on real data, two real datasets were assessed for the presence of interacting SNP pairs.

#### 3.2.1 Rheumatoid arthritis data

After pre-processing and filtering, the Rheumatoid arthritis (RA) dataset contained 94,486 SNPs (4.46 billion possible pairwise interaction tests) and 4,798 individuals. The method, after building a Random Forests analysis, selected only 654 SNP pairs for the second stage of testing to identify significant pairwise interactions (Equation 2). Among the 654 SNP pairs, 26 pairs were found significant using a Bonferroni corrected *p*-value (*p* < 0.05). Specifically, a total of 24 SNP were involved in 26 significant interactions among which 22 of these SNP were from chromosome 6 with two markers from chromosome 1 and chromosome 7. Furthermore, several SNPs from chromosome 6 were involved in multiple interactions. All 22 SNP on chromosome 6 were located in the HLA (human leukocyte antigen) region spanning from 31.35 MB to 32.80 MB. The HLA region in the human genome is known to be highly conserved with high linkage disequilibrium between markers in the region. If interacting SNP pairs are in the same haplotype or LD block, the significant interaction between the pairs might be a result of a haplotype effect on RA, rather than true interaction. Out of the 26 pairs of interacting SNPs, 12 pairs (all markers on chromosome 6) have moderate to high LD (D′ = (0.26 to 0.55)) between the pairs. These were deemed to likely be significant due to being part of the same haplotype. Thus, only 14 SNP pairs with low LD (D′ ≤ 0.2) were retained for further investigation.

Table 4 lists the 14 interacting SNP pairs that were found using the method to be associated with rheumatoid arthritis in humans. Genes containing any of the interacting SNPs as well as flanking genes for intergenic SNP are listed in Supplementary Table S1. All 14 significant SNP pair interactions had at least one SNP of each pair positioned in the major histocompatibility complex (MHC) class II centred on the HLA-DQ to HLA-DR regions located in chromosome 6p21. The HLA region in human chromosome is a 3.78 Mb region that is the most gene dense region within the human genome, encoding about 253 genes including several key immune response genes (Shiina et al., 2009). The genes in this region predominantly have immune response functions. The dominance of the HLA-DQ to HLA-DR region in the interacting SNPs may be due to several factors including the clustering of genes with key roles in immune response and its regulation, high gene density including some overlapping and anti-sense genes, and a high rate of polymorphism and therefore high linkage disequilibrium.

**Table 4.** Fourteen pairs of interacting SNPs have significant odds ratio after pairwise LD evaluation

| SNP pair | D' | r<sup>2</sup> | SNP1 location | SNP1 odds ratio (95% CI)<sup>b</sup> | SNP1 p-value<sup>d</sup> | SNP2 location | SNP2 odds ratio (95% CI)<sup>b</sup> | SNP2 p-value<sup>d</sup> | Interaction odds ratio (95% CI)<sup>c</sup> | Interaction p-value<sup>d</sup> |
| --- | --- | --- | --- | --- | --- | --- | --- | --- | --- | --- |
| rs3134926:rs910050 | 0.10 | 0.01 | 6:32232370 | 0.61 (0.55,0.67) | 1.99E-20 | 6:32347877 | 0.79 (0.72,0.86) | 7.11E-05 | 1.77 (1.53,2.06) | 3.38E-11 |
| rs3134926:rs4530903 | 0.14 | 0.01 | 6:32232370 | 0.61 (0.55,0.67) | 1.99E-20 | 6:32614112 | 1.41 (1.24,1.59) | 6.59E-05 | 1.64 (1.34,2.01) | 9.90E-04 |
| rs9431614:rs9296009 | 0.01 | 0.00 | 1:230944623 | 1.00 (0.89,1.11) | 5.17E+02 | 6:32146738 | 1.80 (1.64,1.97) | 2.51E-33 | 1.62 (1.35,1.96) | 2.91E-04 |
| rs3134926:rs9391858 | 0.06 | 0.00 | 6:32232370 | 0.61 (0.55,0.67) | 1.99E-20 | 6:32373621 | 1.24 (1.11,1.39) | 5.76E-02 | 1.55 (1.28,1.88) | 4.22E-03 |
| rs10456058:rs9273363<sup>a</sup> | 0.08 | 0.00 | 6:31526961 | 1.11 (1.03,1.21) | 5.97E+00 | 6:32658495 | 1.34 (1.23,1.46) | 3.39E-08 | 1.50 (1.32,1.7) | 2.73E-07 |
| rs910050:rs6457617 | 0.17 | 0.02 | 6:32347877 | 0.79 (0.72,0.86) | 7.11E-05 | 6:32696074 | 0.44 (0.4,0.48) | 1.06E-69 | 1.39 (1.22,1.59) | 4.32E-04 |
| rs9268831:rs615672 | 0.15 | 0.01 | 6:32459971 | 0.69 (0.63,0.75) | 3.37E-16 | 6:32606394 | 0.60 (0.54,0.66) | 7.78E-24 | 1.34 (1.17,1.53) | 9.12E-03 |
| rs2076533:rs12706898 | 0.03 | 0.00 | 6:32395750 | 0.49 (0.45,0.54) | 4.60E-55 | 7:129615156 | 1.06 (0.98,1.16) | 8.23E+01 | 1.31 (1.16,1.48) | 1.48E-02 |
| rs17421624:rs615672 | 0.18 | 0.01 | 6:32098400 | 1.70 (1.56,1.86) | 7.24E-31 | 6:32606394 | 0.60 (0.54,0.66) | 7.78E-24 | 0.73 (0.64,0.84) | 7.58E-03 |
| rs6457617:rs2621382 | 0.17 | 0.03 | 6:32696074 | 0.44 (0.4,0.48) | 1.06E-69 | 6:32792668 | 1.02 (0.94,1.11) | 3.44E+02 | 0.73 (0.64,0.83) | 6.46E-04 |
| rs17421624:rs3134926 | 0.08 | 0.00 | 6:32098400 | 1.70 (1.56,1.86) | 7.24E-31 | 6:32232370 | 0.61 (0.55,0.67) | 1.99E-20 | 0.72 (0.62,0.83) | 7.12E-03 |
| rs17421624:rs427037 | 0.03 | 0.00 | 6:32098400 | 1.70 (1.56,1.86) | 7.24E-31 | 6:32244487 | 0.70 (0.62,0.78) | 2.70E-07 | 0.69 (0.59,0.82) | 1.11E-02 |
| rs9296009:rs4530903 | 0.10 | 0.00 | 6:32146738 | 1.80 (1.64,1.97) | 2.51E-33 | 6:32614112 | 1.41 (1.24,1.59) | 6.59E-05 | 0.67 (0.55,0.81) | 3.16E-02 |
| rs206015:rs3134926 | 0.06 | 0.00 | 6:32214982 | 1.49 (1.33,1.67) | 2.90E-09 | 6:32232370 | 0.61 (0.55,0.67) | 1.99E-20 | 0.67 (0.55,0.82) | 4.66E-02 |

<sup>a</sup>rs10456058 has merged into rs2734573.
<sup>b</sup>Odds ratio (OR) of having at least one copy of the risk allele compared to having 0 copies of the same allele for the given SNP.
<sup>c</sup>Odds ratio (OR) of having at least one copy of each risk allele compared to having 0 copies of the same allele for the given SNP pair.
<sup>d</sup>Bonferroni adjusted p-value.

The identification of 12 pairs with both SNP in the HLA-DQ - HLA-DR region on chromosome 6p21 demonstrates that the proposed method is able to identify cis-acting positive and negative SNP interactions identifying potential gene interactions in this gene-dense HLA-DQ - HLA-DR region.

The identified SNPs may be genetic markers closely linked to causal genetic variants or it is also possible that some of the interacting SNPs may be causal. Two identified interacting pairs of SNP (rs615672/rs9268831 and rs92688831/rs615672) were found to have both markers in potential genome regulatory elements. Collectively, this information suggests that local gene-gene interactions in this region could be mediated by cis-acting regulatory mechanisms involving short and long range DNA loops. Some of the identified interactive SNPs were located near or within introns of genes unrelated to the HLA-DQ and HLA-DR genes but interspersed within the same region on chromosome 6p21. These genes included *NOTCH4* (notch receptor 4), *TNXB* (tenascin XB) and *BTNL2* (butryophilin-like 2). *NOTCH4* regulates cell fate determination, proliferation and apoptosis programs and has a regulatory role in branching morphogenesis in the developing vascular system. It is also implicated in the regulation of inflammation (Harb et al., 2020; Zheng et al., (2018), a characteristic feature of autoimmune diseases such as rheumatoid arthritis. *TNXB* encodes a tenascin family member of extracellular matrix proteins. It has anti-adhesive effects and is involved in extracellular matrix maturation following wound healing (Stelzer et al., 2016). Thus, the protein encoded by *TNXB* may potentially be involved in tissue repair after damage caused by chronic inflammation. *BTNL2* belongs to a family of immune regulators. It decreases T-cell proliferation and cytokine release during an inflammatory response and has been implicated in a number autoimmune disease (Arnett et al., 2007; Stelzer et al., 2016; Lebrero-Ferna ndez et al., 2016). The SNP interaction partner to the SNP identified in *BTNL2* on 6p21 is located near *NRF1* (nuclear respiratory facto 1) on chromosome 7. *NRF1* encodes a transcription factor activating key metabolic and cell growth genes as well as mitochondrial DNA transcription and replication. *NRF1* has also been implicated in the regulation of inflammation, Stelzer et al. (2016); van Tienen et al. (2010); Yang et al. (2018). One possibility is that *NRF1* acts in trans directly or indirectly via its encoded transcription factor to regulate *BTNL2* expression on chromosome 6p21 and thus modulate T-cell mediated immune activation and inflammation.

Notably, multiple genome wide association studies (GWAS) identified a subset of the interactive SNPs as associated with various autoimmune diseases i.e., rs9273363, rheumatoid arthritis; type 1 diabetes; rs6457617, multiple sclerosis, Graves disease, systemic sclerosis; rs615672, rheumatoid arthritis (“UCSC Genome Browser Home”).

The results from using the proposed method on the human rheumatoid arthritis data revealed a number of promising results supporting the potential value of the proposed method for identifying epistatic interactions in case control data including the ability to find positive and negative cis-acting interactions.

#### 3.2.2 Mouse data

The curated mouse data contained 9,265 SNPs resulting in nearly 43 million possible pairwise comparisons to test for the presence of significant interactions contributing to the phenotype “weight at 6 weeks”. The proposed method, identified 273,391 pairs of SNP (0.0064%) for the second stage of interaction testing. After using the Bonferroni multiple testing correction, 59 pairs of interacting SNPs remained as significant. These 59 pairs of interactions were from 63 unique SNPs mainly located on MMU chromosomes 2, 8 and 13. To check the validity of the results and proposed method, the genes associated with each of these SNP were identified using the Mouse Genome Informatics (The Jackson Laboratory, 2021). Supplementary Table S2 lists the significant interacting pairs of SNPs that were found associated with the murine growth trait W6W. The nearest genes (mean distance = 36.7 Kbp; range = 42 bp – 286 Kbp) associated with these SNP are also listed. The associated MGI ID (Mouse Genome Informatics ID), Entrez ID and GO terms (Gene Ontology) are listed in Supplementary Table S3.

The proposed method identified four genes and two functionally anonymous genes (*miR8108* and *Gm31784*) associated with two additional SNPs, in a 3.8 Mbp interval on MMU Chr8qA2. Genes *Tm2d2, Tacc1, Slc20a2* and *Fgfr1* are contained within a small region of 0.5 Mbp of this larger segment and have a variety of functions that potentially impact a phenotype associated with growth and development. *Fgfr1* (fibroblast growth factor receptor 1) has wide ranging biological functions associated with regulation of cell division, cell growth and maturation, embryonic development, wound healing, and formation of blood vessels. In humans, the cellular signalling mediated by this receptor has roles in the development and growth of bones, particularly bones in the head, face, hand, feet, and long bones of the arms and legs. Mutations in the gene are associated with human dwarfism (National Institutes of Health, 2021). *Tacc1* (transforming acidic coiled-coil containing protein 1) is involved in processes that promote cell division before the formation of differentiated tissues (Stelzer et al. 2016). *Tacc1* encodes a protein that regulates the localization of nuclear receptors, including T3 thyroid hormone and retinoic pathways involved in regulation cell growth and differentiation. *Tm2d2* (TM2 domain containing 2) encodes a G-coupled receptor that regulates cell death and proliferation cascades (Stelzer et al., 2016). *Slc20a2* (solute carrier family 20 member 2) encodes an inorganic phosphate transporter important in maintaining phosphate homeostasis. This function is essential for all intracellular function. The encoded protein also has roles in extracellular matrix and cartilage calcification, which is important for tissue function (Stelzer et al., 2016). It is noted that growing bones have high demands for phosphate (Goretti Penido and Alon 2012).

Four genes identified as potentially involved in interactions were located in a 0.25 Mbp region on the MMU Chr2qA3-2qB boundary. They were (*Mrps2* (mammalian mitochondrial ribosomal protein), *Ralgds* (ral guanine nucleotide dissociation stimulator), *Ddx31* (dead-box helicase 31) and *Ttf1* (transcription termination factor 1). These genes have a variety of functions including mitochondrial protein synthesis, intracellular signalling, cell growth and division in embryogenesis, and transcription, respectively (Stelzer et al., 2016). At present, only *Ddx31* is directly linked to growth related traits, although the molecular functions of the remaining three genes may have this potential. The remaining genes identified from the analyses, listed in Tables S2 and S3, map to various chromosomal regions. Notable gene functions or mutational effects that could influence growth traits include: regulation of cell proliferation, differentiation and survival (*Klf7*, Kruppel like factor 7); osteogenesis (*Stmn2*, Stathmin 2); short stature and skeletal abnormalities (*Wipi2*, WD40 repeat protein interacting with phosphoinositides 2); increased blood testosterone, precocious sexual maturation, increased fecundity, and clearance of luteinizing hormone (*Chst8*, carbohydrate sulfo-transferase 8) (Stelzer et al. 2016)). In all, there were several genes identified close to interacting SNPs listed in Tables S2 and S3 that have effects on murine growth and developmental traits, particularly osteogenesis and body stature.

Analysis of the mouse data results revealed strong gene candidates for involvement in growth and development pathways in mice, including genes known to have relevant biological function as well as novel candidates. This finding indicates that the function of these genes could be influenced by interactions with other genes and that the proposed method is able to identify multiple promising regions for a quantitative trait for further exploration.

## Conclusion

In this study, a novel efficient method to identify potentially interacting pairs of markers in genomic data for a specific trait was proposed and validated. A Random Forests decision tree algorithm was utilised to select a subset of markers pairs from, the often, vast search space to enable the efficient computational testing of these pairs for evidence of significant interaction. The method was validated with several simulation datasets representing different data structures and characteristics including marker density and trait heritability. The method was then used with two real data sets to identify SNP interactions for two different type of traits, case-control data for the presence of rheumatoid arthritis in humans, and quantitative trait data for the body weight of mice. In both cases, the method identified several functionally relevant genes positioned close to the markers shown to be potentially interacting for the respective traits. The method is demonstrated as having potential in providing new insights into the genetic basis of complex traits controlled by multiple genes and their interactions. The method can be extended in a biological context beyond purely genomic data to other ’omic data types or interactions between diverse types of data such as marker and environmental data to further provide biological insights.

## Key points

- Detecting epistatic interactions from high dimensional genomic data is very challenging due to the computational burden induced by the large number of required combinatorial test.
- Using a Random Forests algorithm, we provided a computationally simple way to detect pairwise interactions.
- The method produced reliable results in the case of simulated and interpretable results in several real-world data sets.

## Data availability

PIC nucleus pig line genotype data is available as supporting information in Cleveland et al., 2012. WTCCC genotype data is available by application to the Wellcome Trust Case Control Consortium Data Access Committee. The mouse dataset is available as Additional file 1 in (Martini et al., 2017).

## Author contributions statement

H.A., R.D and K.V. conceived the experiment(s), H.A. conducted the experiment(s), H.A., R.D. and R.T. analysed the results. H.A., R.D., K.V. and R.T. wrote and reviewed the manuscript.

## Acknowledgements

The authors thank the anonymous reviewers for their valuable suggestions. This work is supported by CSIRO MLAI Future Science Platform. This study makes use of data generated by the Wellcome Trust Case Control Consortium. A full list of the investigators who contributed to the generation of the data is available from www.wtccc.org.uk. Funding for the project was provided by the Wellcome Trust under award 076113.

## References

Ansarifar, J., Akhavizadegan, F., and Wang, L. (2020). Performance prediction of crosses in plant breeding through genotype by environment interactions. Sci Rep, 10(1):11533.

Arnett, H. A., Escobar, S. S., Gonzalez-Suarez, E., Budelsky, A. L., Steffen, L. A., Boiani, N., Zhang, M., Siu, G., Brewer, A. W., and Viney, J. L. (2007). BTNL2, a Butyrophilin/B7-Like Molecule, Is a Negative Costimulatory Molecule Modulated in Intestinal Inflammation. The Journal of Immunology, 178(3):1523–1533.

Auton, A. and more than 93 other authors (2015). A global reference for human genetic variation. Nature, 526(7571):68– 74.

Breiman, L. (2001). Random Forests. Machine Learning, 45(1):5–32.

Burton, P. R. and 297 other authors (2007). Genome-wide association study of 14,000 cases of seven common diseases and 3,000 shared controls. Nature, 447(7145):661–678.

Cleveland, M. A., Hickey, J. M., and Forni, S. (2012). A Common Dataset for Genomic Analysis of Livestock Populations. G3: Genes— Genomes—Genetics, 2(4):429– 435.

Gholami, M., Erbe, M., Garke, C., Preisinger, R., Weigend, A., Weigend, S., and Simianer, H. (2014). Population genomic analyses based on 1 million snps in commercial egg layers. PLoS One, 9(4):e94509.

Goretti Penido, M. and Alon, U. S. (2012). Phosphate homeostasis and its role in bone health. Pediatr Nephrol, 27(11):2039–2048.

Ha, N. T., Freytag, S., and Bickeboeller, H. (2014). Coverage and efficiency in current snp chips. Eur J Hum Genet, 22(9):1124–30.

Harb, H., Stephen-Victor, E., Crestani, E., Benamar, M., Massoud, A., Cui, Y., Charbonnier, L. M., Arbag, S., Baris, S., Cunnigham, A., Leyva-Castillo, J. M., Geha, R. S., Mousavi, A. J., Guennewig, B., Schmitz-Abe, K., Sioutas, C., Phipatanakul, W., and Chatila, T. A. (2020). A regulatory t cell notch4-gdf15 axis licenses tissue inflammation in asthma. Nat Immunol, 21(11):1359–1370.

Jiang, R., Tang, W., Wu, X., and Fu, W. (2009). A Random Forests approach to the detection of epistatic interactions in case-control studies. BMC Bioinformatics, 10 Suppl 1:S65.

Lebrero-Fernandez, C., Wenzel, U. A., Akeus, P., Wang, Y., Strid, H., Simr en, M., Gustavsson, B., Bo □rjesson, L. G., Cardell, S. L., O □hman, L., Quiding-J □arbrink, M., and Bas-Forsberg, A. (2016). Altered expression of Butyrophilin (BTN) and BTN-like (BTNL) genes in intestinal inflammation and colon cancer. Immunity, Inflammation and Disease, 4(2):191–200.

Lundberg, S. M., Erion, G. G., and Lee, S.-I. (2019). Consistent Individualized Feature Attribution for Tree Ensembles. arXiv:1802.03888 [cs, stat].

Maher, B. (2008). Personal genomes: The case of the missing heritability. Nature, 456(7218):18–21.

Martini, J. W. R., Gao, N., Cardoso, D. F., Wimmer, V., Erbe, M., Cantet, R. J. C., and Simianer, H. (2017). Genomic prediction with epistasis models: On the markercoding-dependent performance of the extended GBLUP and properties of the categorical epistasis model (CE). BMC Bioinformatics, 18(1):3.

Meng, Y., Yang, Q., Cuenco, K. T., Cupples, L. A., DeStefano, A. L., and Lunetta, K. L. (2007). Two-stage approach for identifying singlenucleotide polymorphisms associated with rheumatoid arthritis using Random Forests and bayesian networks. BMC Proceedings, 1(Suppl 1):S56.

Meng, Y. A., Yu, Y., Cupples, L. A., Farrer, L. A., and Lunetta, K. L. (2009). Performance of Random Forests when SNPs are in linkage disequilibrium. BMC Bioinformatics, 10(1):78.

National Institutes of Health (2021). Medline Plus, National Institutes of Health, U.S. National Library of Medicine. https://medlineplus.gov/genetics. Accessed: Monday, May 10, 2021.

Pecanka, J., Jonker, M. A., International Parkinson, S. D. G. C., Bochdanovits, Z., and Van Der Vaart, A. W. (2017). A powerful and efficient two-stage method for detecting geneto-gene interactions in GWAS. Biostatistics, 18(3):477–494.

Purcell, S., Neale, B., Todd-Brown, K., Thomas, L., Ferreira, M. A. R., Bender, D., Maller, J., Sklar, P., de Bakker, P. I. W., Daly, M. J., and Sham, P. C. (2007). PLINK: A Tool Set for Whole-Genome Association and Population-Based Linkage Analyses. American Journal of Human Genetics, 81(3):559–575.

Schmalohr, C. L., Grossbach, J., Clement-Ziza, M., and Beyer, A. (2018). Detection of epistatic interactions with Random Forests. bioRxiv, page 353193.

Shiina, T., Hosomichi, K., Inoko, H., and Kulski, J. K. (2009). The HLA genomic loci map: Expression, interaction, diversity and disease. Journal of Human Genetics, 54(1):15– 39.

Solberg, L. C., Valdar, W., Gauguier, D., Nunez, G., Taylor, A., Burnett, S., Arboledas-Hita, C., Hernandez-Pliego, P., Davidson, S., Burns, P., Bhattacharya, S., Hough, T., Higgs, D., Klenerman, P., Cookson, W. O., Zhang, Y., Deacon, R. M., Rawlins, J. N. P., Mott, R., and Flint, J. (2006). A protocol for high-throughput phenotyping, suitable for quantitative trait analysis in mice. Mammalian Genome: Official Journal of the International Mammalian Genome Society, 17(2):129– 146.

Stelzer, G., Rosen, N., Plaschkes, I., Zimmerman, S., Twik, M., Fishilevich, S., Stein, T. I., Nudel, R., Lieder, I., Mazor, Y., Kaplan, S., Dahary, D., Warshawsky, D., Guan-Golan, Y., Kohn, A., Rappaport, N., Safran, M., and Lancet, D. (2016). The genecards suite: From gene data mining to disease genome sequence analyses. Current Protocols in Bioinformatics, 54(1):1.30.1–1.30.33.

The Jackson Laboratory, Bar Harbor, M. (2021). Mouse genome informatics. http://www.informatics.jax.org/. Accessed: Monday, May 10, 2021.

University of Utah (2021). Whole Genome SNP Genotyping. https://cores.utah.edu/genomics/ genomics-whole-genome-snpgenotyping. Accessed: Monday, May 10, 2021.

Valdar, W., Solberg, L. C., Gauguier, D., Cookson, W. O., Rawlins, J. N. P., Mott, R., and Flint, J. (2006). Genetic and environmental effects on complex traits in mice. Genetics, 174(2):959–984.

van Tienen, F. H., Lindsey, P. J., van der Kallen, C. J., and Smeets, H. J. (2010). Prolonged Nrf1 overexpression triggers adipocyte inflammation and insulin resistance. Journal of Cellular Biochemistry, 111(6):1575–1585.

Wright, M. N. and Ziegler, A. (2017). ranger: A fast implementation of Random Forests for high dimensional data in C++ and R. Journal of Statistical Software, 77(1):1–17.

Yang, K., Huang, R., Fujihira, H., Suzuki, T., and Yan, N. (2018). N-glycanase NGLY1 regulates mitochondrial homeostasis and inflammation through NRF1. The Journal of Experimental Medicine, 215(10):2600–2616.

Yoshida, M. and Koike, A. (2011). SNPInterForest: A new method for detecting epistatic interactions. BMC Bioinformatics, 12(1):469.

Zhang, X., Zou, F., and Wang, W. (2008). Fastanova: an efficient algorithm for genome-wide association study. KDD, pages 821–829.

Zheng, R., Liu, H., Zhou, Y., Yan, D., Chen, J., Ma, D., Feng, Y., Qin, L., Liu, F., Huang, X., Wang, J., and Ge, B. (2018). Notch4 negatively regulates the inflammatory response to mycobacterium tuberculosis infection by inhibiting tak1 activation. J Infect Dis, 218(2):312–323.

Ziegler, A., DeStefano, A. L., Konig, I. R., Bardel, C., Brinza, D., Bull, S., Cai, Z., Glaser, B., Jiang, W., Lee, K. E., Li, C. X., Li, J., Li, X., Majoram, P., Meng, Y., Nicodemus, K. K., Platt, A., Schwarz, D. F., Shi, W., Shugart, Y. Y., Stassen, H. H., Sun, Y. V., Won, S., Wang, W., Wahba, G., Zagaar, U. A., and Zhao, Z. (2007). Data mining, neural nets, trees–problems 2 and 3 of genetic analysis workshop 15. Genet Epidemiol, 31 Suppl 1:S51–60.

Zuk, O., Hechter, E., Sunyaev, S. R., and Lander, E. S. (2012). The mystery of missing heritability: Genetic interactions create phantom heritability. Proc Natl Acad Sci USA, 109(4):1193–8.

