## Supplementary Table S1 for "Detecting epistatic interactions in genomic data using Random Forests"

**(A) Interacting SNP pairs for rheumatoid arthritis data**

| **SNP pair** | ***SNP1 location*** | ***SNP2 location*** | **intronic** | **intergenic** | **SNP regulatory** | **GWAS** |
| --- | --- | --- | --- | --- | --- | --- |
| rs3134926:rs910050 | chr6:32232370 | chr6:32347877 | SNP2, intronic in C6orf10 (TSBP1) | SNP1 intergenic between NOTCH4 and TSBP1 | SNP1, nil; SNP2, nil |  |
| rs3134926:rs4530903 | chr6:32232370 | chr6:32614112 |  | SNP1 intergenic between NOTCH4 and TSBP1; SNP2 between HLA-DRB1 and HLA-DQA1 | SNP1 nil; SNP2, nil | SNP2, coronary heart disease; endometriosis; SNP2, lymphoma, non-Hodgkins lymphoma, schizophrenia |
| rs9431614:rs9296009 | chr1:230944623 | chr6:32146738 | SNP1 intronic in TTC13 on chr 1 | SNP2 intergenic between FKBPL and PRRT1 on chr 6 | SNP1 nil; SNP2 nil | SNP2, blood protein levels, IgG glycosylation |
| rs3134926:rs9391858 | chr6:32232370 | chr6:32373621 | SNP2, intronic in TSBP1_AS1 | SNP1 intergenic between NOTCH4 and TSBP1 | SNP1, nil; SNP2, nil | SNP2, total cholesterol levels |
| *rs10456058 (rs2734573):rs9273363 | chr6:31526961 | chr6:32658495 |  | SNP1 intergenic between MICB and MCCD1; SNP2 between HLA-DQA1 and HLA-DQB1 and HLA-DQB1-AS1 | SNP1, nil; SNP2, nil | SNP2, type 1 diabetes risk, chronic lymphocytic leulemia |
| rs910050:rs6457617 | chr6:32347877 | chr6:32696074 | SNP1, intronic in C6orf10 (TSBP1) and TSBP1_AS1 | SNP2 between HLA-DQB1 and HLA-DQA2 | SNP1, distal enhancer; SNP2, nil | SNP2, thyrotoxic hypokalemic paralysis, IgG index levels, Graves Disease, multiple sclerosis, rheumatoid arthritis. |
| rs9268831:rs615672 | chr6:32459971 | chr6:32606394 |  | SNP1 between HLA-DRA and HLA-DRB5; SNP2, between HLA-DRB6 and HLA-DQA1; | SNP1 distal enhancer | SNP1, response to hepatitis vaccine; SNP2 waist cirumference adjusted for BMI, physical activity interactions in adiposity; rheumatoid arthritis (Nature 2007 Wellcome Trust Case Control Consortium; |
| rs2076533:rs12706898 | chr6:32395750 | chr7:129615156 | SNP1 intronic in TSBP1_AS1 and BTNL2; SNP2 intronic in NRF1 |  | SNP1 nil; SNP2 distal enhancer |  |
| rs17421624:rs615672 | chr6:32098400 | chr6:32606394 | SNP1 intronic in TNXB; | SNP2, between HLA-DRB6 and HLA-DQA1; | SNP1 nil; SNP2 nil | SNP2 waist cirumference adjusted for BMI, physical activity interactions in adiposity; rheumatoid arthritis (Nature 2007 Wellcome Trust Case Control Consortium; |
| rs6457617:rs2621382 | chr6:32696074 | chr6:32792668 |  | SNP1 intergenic between HLA-DQB1 and HLA-DQA2 | SNP2, distal enhancer | SNP1, thyrotoxic hypokalemic paralysis, IgG index levels, Graves Disease, multiple sclerosis, rheumatoid arthritis. |
| rs17421624:rs3134926 | chr6:32098400 | chr6:32232370 | SNP1 intronic in TNXB; SNP2 intronic in TSBP1_AS1 and BTNL2 | SNP2 intergenic between NOTCH4 and TSBP1 | SNP1 nil; SNP2 nil |  |
| rs17421624:rs427037 | chr6:32098400 | chr6:32244487 | SNP1 intronic in TNXB; | SNP2 intergenic between NOTCH4 and TSBP1-AS1 | SNP1 nil; SNP distal enhancer |  |
| rs9296009:rs4530903 | chr6:32146738 | chr6:32614112 |  | SNP1 intergenic between FKBPL and PRRT1 on chr 6; intergenic between HLA-DRB1 and HLA-DQA1 | SNP1 nil; SNP2 nil | SNP1, blood protein levels, IgG glycosylation; SNP2 lymphoma, schizophrenia |
| rs206015:rs3134926 | chr6:32214982 | chr6:32232370 | SNP1 intronic in NOTCH4 | ; SNP2 intergenic between NOTCH4 and TSBP1 | SNP1 nil; SNP2 nil |  |

*^*^*rs10456058 has merged into rs2734573.

**(B) Genes and flanking genes**

| ***Gene Symbol*** | **Gene name** | **comment** | **inflammation** | **Associated withautoimmune diseases** |
| --- | --- | --- | --- | --- |
| *NOTCH4* | NOTCH neurogenic gene 4 | role in vascular development; cell fate determination | negatively regulates inflammation | PMID: 32929274; PMID: 31838262; PMID: 30442821; PMID: 29693251; PMID: 29228365; PMID: 33134369 |
| *TSBP1* | testis basic protein 1 | seems irrelevant to RA; highly testes specific expression | | |
| *TSBP1-AS1* | antisense to TSBP1 |  |  |  |
| *C6orf10 (TSBP1)* | same as TSBP1 | seems irrelevant to RA; highly testes specific expression | | |
| *BTNL2* | butyrophilin-like B7 | immunoregulator and surveilance; negative T-cell regulator; decreases T-cell proliferation and cytokine release | negative regulator of inflammation | rhematoid arthritis; sucroidosis; ulcerative colitis; intestinal bowel disease; myositis; type 1 diabetes; systemic lupis erythematous; acute coronary syndrome; prostate cancer |
| *MICB* | MHC class I polypeptide-related sequence B | ligand for NKG2D type II receptor; role in activation of cytolytic response of natural killer cells, CD8 alpha beta cells, gamma delta T cells; stress induced | positively regulates inflammation | |
| *MCCD1* | mitochondrial coiled coil domain 1 | nuclear gene for mitochondrial product; overlaps TSBP1_AS1 | | |
| *TTC13* | tetratricopeptide family of extracellular matrix glycoproteins | anti-adhesive effects; matrix maturation during wound healing; Ehlers-Danios syndrome (hyper limb mobility) | | |
| *FKBPL* | proline isomerase | immunophilin family; immunoregulation | binds immunosuppressants | |
| *PRRT1* | prolin rich transmembrane protein | dispanin family; not much known about this gene | | |
| *TNXB* | tenascin XB | tenascin family of extracellulat matrix proteins; anti adhesive; ECM maturation and wound healing; Ehlers-Danlos hyper limb mobility; overlaps with CYP21A2 (cytochrome P450 family subfamily A member 2)) | Woumd healing therefore anti-inflammatory | |
| *C2* | complement C2 | class III MHC region; serum glycoprotein; part of complement classical pathway; blood clotting; | blood clotting | C2 defiency associated with autoimmune diseaeses; SNPs in gene assoc with macular degeneration |
| *CFAP54* | cilia and flagella associated protein 54 | very little known of function | | associated with amyotrophic lateral sclerosis |
| *NEDD1* | gamma-tubulin ring complex factor | cell division |  |  |
| *SKIV2L* | ski2-like RNA helicase | not much known |  | age related macular degeneration; ??tricho-hepato-enteric syndrome |
| *GPANK1* | G-patch domain and ankyrin repeats 1 | thought to play a role in immunity; not much known | PMID: 28282431 ?? | associated with non-Hodhins lymphoma; early menopause; celiac disease |
| *MICA* | MHC class I polypeptide-related sequence A | highly polymorphic; stress induced antigen activation of gamma delta T cells; | T cell activation | psorasis, psoriatic arthritis |
| *MICA-AS1* | antisense to MICA |  |  |  |
| *NRF1* | nuclear respiratory factor 1 | activates key metabolic genes during cell growth and genes required for respiration, mitachondrial DNA transcription and replication; neurite outgrowth | | |
| *HLA-DRA* | major histocompatability complex, class II, DRA | central role in immune system; presents foreign (and self during development) peptides; expressed by antigen presenting cells. Elicits or suppresses T(helper) cell responses that eventually lead to production of antibody against potential antigen. | inflammation |  |
| *HLA-DRB* | major histocompatability complex, class II, DRB | central role in immune system; presents foreign (and self during development) peptides; expressed by antigen presenting cells. Elicits or suppresses T(helper) cell responses that eventually lead to production of antibody against potential antigen. | inflammation |  |
| *HLA-DRB6* | major histocompatability complex, class II, DRB6 | central role in immune system; presents foreign (and self during development) peptides; expressed by antigen presenting cells. Elicits or suppresses T(helper) cell responses that eventually lead to production of antibody against potential antigen. | inflammation |  |
| *HLA-DQA1* | major histocompatability complex, class II, DQA1 | central role in immune system; presents foreign (and self during development) peptides; expressed by antigen presenting cells. Elicits or suppresses T(helper) cell responses that eventually lead to production of antibody against potential antigen. | inflammation |  |
| *HLA-DQA2* | major histocompatability complex, class II, DQA2 | central role in immune system; presents foreign (and self during development) peptides; expressed by antigen presenting cells. Elicits or suppresses T(helper) cell responses that eventually lead to production of antibody against potential antigen. | inflammation |  |
| *HLA-DQB1* | major histocompatability complex, class II, DQB1 | central role in immune system; presents foreign (and self during development) peptides; expressed by antigen presenting cells. Elicits or suppresses T(helper) cell responses that eventually lead to production of antibody against potential antigen. | inflammation |  |
| *HLA-DQB1-AS1* | major histocompatability complex, class II, DQB!-AS1 | central role in immune system; presents foreign (and self during development) peptides; expressed by antigen presenting cells. Elicits or suppresses T(helper) cell responses that eventually lead to production of antibody against potential antigen. | inflammation |  |
| *HLA-DRB1* | major histocompatability complex, class II, DRB1 | central role in immune system; presents foreign (and self during development) peptides; expressed by antigen presenting cells. Elicits or suppresses T(helper) cell responses that eventually lead to production of antibody against potential antigen. | inflammation |  |
| *HLA-DRB5* | major histocompatability complex, class II, DRB5 | central role in immune system; presents foreign (and self during development) peptides; expressed by antigen presenting cells. Elicits or suppresses T(helper) cell responses that eventually lead to production of antibody against potential antigen. | inflammation |  |
| *HLA-DRB6* | major histocompatability complex, class II, DRB6 | central role in immune system; presents foreign (and self during development) peptides; expressed by antigen presenting cells. Elicits or suppresses T(helper) cell responses that eventually lead to production of antibody against potential antigen. | inflammation |  |
| *HLA-B* | major histocompatability complex, class II, B | central role in immune system; presents foreign (and self during development) peptides; expressed by antigen presenting cells. Elicits or suppresses T(helper) cell responses that eventually lead to production of antibody against potential antigen. | inflammation |  |
