## Supplementary Table S2 for "Detecting epistatic interactions in genomic data using Random Forests"

**Supplementary Table S2**: List of the significant interacting pairs of SNPs associated with the murine growth trait W6W.

| **SNP1** | **SNP2** | **SNP1**  **effect** | **SNP1**  **P-value** | **SNP2**  **effect** | **SNP2**  **P-value** | **Interaction**  **effect** | **Interaction**  **P-value** | **SNP1 Mbp** | **SNP1**  **CHR** | **SNP2 Mbp** | **SNP2**  **CHR** | **SNP1**  **Nearest**  **gene** | **SNP1**  **Nearest gene**  **Start pos** | **SNP1**  **Nearest gene end pos** | **SNP1**  **Nearest gene distance in bp** | **SNP1 Nearest gene description** | **SNP2**  **Nearest**  **gene** | **SNP2 Nearest gene start pos** | **SNP2 Nearest gene end pos** | **SNP2 Nearest gene distance in bp** | **SNP2 Nearest gene description** |
| --- | --- | --- | --- | --- | --- | --- | --- | --- | --- | --- | --- | --- | --- | --- | --- | --- | --- | --- | --- | --- | --- |
| rs3163007 | rs13459102 | -0.002 | 0.63432056 | -0.099 | 1.60E-06 | 0.106 | 0.00398038 | 64.041 | 1 | 25.023 | 8 | Mir6899 | 64042438 | 64042501 | 1359.5 |  | Tm2d2 | 25017211 | 25023260 | 2432.5 | TM2 domain containing 2 |
| rs3163007 | rs13479654 | -0.002 | 0.6238091 | -0.101 | 1.05E-06 | 0.109 | 0.00172817 | 64.041 | 1 | 25.178 | 8 | Mir6899 | 64042438 | 64042501 | 1359.5 |  | Mir8108 | 25159125 | 25159233 | 18936 |  |
| rs3162895 | rs13479654 | -0.002 | 0.67363584 | -0.101 | 1.13E-06 | 0.109 | 0.00196586 | 64.201 | 1 | 25.178 | 8 | Gm28981 | 64185991 | 64303043 | 43076 |  | Mir8108 | 25159125 | 25159233 | 18936 |  |
| rs3163007 | rs13479656 | -0.002 | 0.62596873 | -0.094 | 4.84E-06 | 0.105 | 0.00574361 | 64.041 | 1 | 25.532 | 8 | Mir6899 | 64042438 | 64042501 | 1359.5 |  | Fgfr1 | 25513654 | 25575718 | 12261 | fibroblast growth factor receptor 1 |
| rs3162895 | rs3704385 | -0.002 | 0.67558972 | -0.093 | 5.10E-06 | 0.105 | 0.00651667 | 64.201 | 1 | 26.305 | 8 | Gm28981 | 64185991 | 64303043 | 43076 |  | Gm31784 | 26298352 | 26312334 | 582 |  |
| rs6212146 | rs3704385 | -0.002 | 0.72432031 | -0.093 | 5.44E-06 | 0.105 | 0.00733376 | 64.407 | 1 | 26.305 | 8 | Gm25748 | 64396334 | 64396403 | 11129.5 |  | Gm31784 | 26298352 | 26312334 | 582 |  |
| rs13476400 | rs3694031 | 0.010 | 0.37260936 | 0.035 | 1.08E-11 | -0.094 | 0.00378592 | 28.470 | 2 | 34.915 | 7 | Mrps2 | 28468066 | 28471178 | 42 | mitochondrial ribosomal protein S2 | Gm25922 | 34943856 | 34943985 | 29040.5 |  |
| rs13476402 | rs3663861 | 0.001 | 0.9387265 | 0.009 | 0.12677276 | -0.126 | 1.39E-05 | 28.880 | 2 | 69.655 | 9 | Ddx31 | 28840406 | 28905571 | 6546.5 | DEAD/H (Asp-Glu-Ala-Asp/His) box polypeptide 31 | Gm47203 | 69599307 | 69599754 | 55439.5 |  |
| rs3023797 | rs3719311 | 0.003 | 0.75020939 | 0.037 | 1.72E-10 | -0.107 | 0.00130876 | 29.087 | 2 | 34.233 | 7 | Gm13393 | 29088666 | 29094501 | 4767.5 |  | Gm12762 | 34230420 | 34234102 | 802 |  |
| rs13476400 | rs6239372 | 0.009 | 0.37515309 | 0.035 | 9.83E-12 | -0.094 | 0.00366517 | 28.470 | 2 | 34.699 | 7 | Mrps2 | 28468066 | 28471178 | 42 | mitochondrial ribosomal protein S2 | Gm12756 | 34665413 | 34673999 | 29145 |  |
| rs13476402 | rs6239372 | 0.009 | 0.37515309 | 0.035 | 9.83E-12 | -0.094 | 0.00366517 | 28.880 | 2 | 34.699 | 7 | Ddx31 | 28840406 | 28905571 | 6546.5 | DEAD/H (Asp-Glu-Ala-Asp/His) box polypeptide 31 | Gm12756 | 34665413 | 34673999 | 29145 |  |
| rs3023797 | rs6239372 | 0.009 | 0.41179492 | 0.035 | 1.11E-11 | -0.093 | 0.00420837 | 29.087 | 2 | 34.699 | 7 | Gm13393 | 29088666 | 29094501 | 4767.5 |  | Gm12756 | 34665413 | 34673999 | 29145 |  |
| rs13476402 | rs3694031 | 0.010 | 0.37260936 | 0.035 | 1.08E-11 | -0.094 | 0.00378592 | 28.880 | 2 | 34.915 | 7 | Ddx31 | 28840406 | 28905571 | 6546.5 | DEAD/H (Asp-Glu-Ala-Asp/His) box polypeptide 31 | Gm25922 | 34943856 | 34943985 | 29040.5 |  |
| rs6308258 | rs3663861 | 0.001 | 0.9387265 | 0.009 | 0.12677276 | -0.126 | 1.39E-05 | 28.546 | 2 | 69.655 | 9 | Ralgds | 28513125 | 28553081 | 13089 | ral guanine nucleotide dissociation stimulator | Gm47203 | 69599307 | 69599754 | 55439.5 |  |
| rs3023797 | rs3663861 | 0.000 | 0.98343665 | 0.009 | 0.13887351 | -0.120 | 9.24E-05 | 29.087 | 2 | 69.655 | 9 | Gm13393 | 29088666 | 29094501 | 4767.5 |  | Gm47203 | 69599307 | 69599754 | 55439.5 |  |
| rs13476402 | rs3719311 | 0.004 | 0.70237793 | 0.037 | 1.55E-10 | -0.108 | 0.00115679 | 28.880 | 2 | 34.233 | 7 | Ddx31 | 28840406 | 28905571 | 6546.5 | DEAD/H (Asp-Glu-Ala-Asp/His) box polypeptide 31 | Gm12762 | 34230420 | 34234102 | 802 |  |
| rs6308258 | rs3694031 | 0.010 | 0.37260936 | 0.035 | 1.08E-11 | -0.094 | 0.00378592 | 28.546 | 2 | 34.915 | 7 | Ralgds | 28513125 | 28553081 | 13089 | ral guanine nucleotide dissociation stimulator | Gm25922 | 34943856 | 34943985 | 29040.5 |  |
| rs3023797 | rs3694031 | 0.009 | 0.4090622 | 0.035 | 1.22E-11 | -0.093 | 0.00434734 | 29.087 | 2 | 34.915 | 7 | Gm13393 | 29088666 | 29094501 | 4767.5 |  | Gm25922 | 34943856 | 34943985 | 29040.5 |  |
| rs13476400 | rs13479188 | 0.009 | 0.40374323 | 0.034 | 5.12E-11 | -0.093 | 0.00494772 | 28.470 | 2 | 35.340 | 7 | Mrps2 | 28468066 | 28471178 | 42 | mitochondrial ribosomal protein S2 | Rhpn2 | 35334170 | 35392289 | 23391.5 | rhophilin, Rho GTPase binding protein 2 |
| rs13476402 | rs13479188 | 0.009 | 0.40374323 | 0.034 | 5.12E-11 | -0.093 | 0.00494772 | 28.880 | 2 | 35.340 | 7 | Ddx31 | 28840406 | 28905571 | 6546.5 | DEAD/H (Asp-Glu-Ala-Asp/His) box polypeptide 31 | Rhpn2 | 35334170 | 35392289 | 23391.5 | rhophilin, Rho GTPase binding protein 2 |
| rs13476400 | rs3663861 | 0.001 | 0.9387265 | 0.009 | 0.12677276 | -0.126 | 1.39E-05 | 28.470 | 2 | 69.655 | 9 | Mrps2 | 28468066 | 28471178 | 42 | mitochondrial ribosomal protein S2 | Gm47203 | 69599307 | 69599754 | 55439.5 |  |
| rs13469412 | rs3679568 | 0.038 | 8.89E-09 | 0.011 | 0.09084734 | -0.046 | 0.00467236 | 129.134 | 2 | 13.373 | 6 | Chchd5 | 129129700 | 129134134 | 1837 | coiled-coil-helix-coiled-coil-helix domain containing 5 |  | 13413337 | 13415996 | 41895.5 |  |
| rs13476400 | rs3719311 | 0.004 | 0.70237793 | 0.037 | 1.55E-10 | -0.108 | 0.00115679 | 28.470 | 2 | 34.233 | 7 | Mrps2 | 28468066 | 28471178 | 42 | mitochondrial ribosomal protein S2 |  |  |  |  |  |
| rs6308258 | rs3719311 | 0.004 | 0.70237793 | 0.037 | 1.55E-10 | -0.108 | 0.00115679 | 28.546 | 2 | 34.233 | 7 | Ralgds | 28513125 | 28553081 | 13089 | ral guanine nucleotide dissociation stimulator |  |  |  |  |  |
| rs6308258 | rs6239372 | 0.009 | 0.37515309 | 0.035 | 9.83E-12 | -0.094 | 0.00366517 | 28.546 | 2 | 34.699 | 7 | Ralgds | 28513125 | 28553081 | 13089 | ral guanine nucleotide dissociation stimulator |  |  |  |  |  |
| rs6308258 | rs13479188 | 0.009 | 0.40374323 | 0.034 | 5.12E-11 | -0.093 | 0.00494772 | 28.546 | 2 | 35.340 | 7 | Ralgds | 28513125 | 28553081 | 13089 | ral guanine nucleotide dissociation stimulator |  |  |  |  |  |
| rs13476396 | rs3663861 | 0.019 | 0.05460622 | 0.007 | 0.23577062 | -0.131 | 0.00039857 | 27.646 | 2 | 69.655 | 9 | Rxra | 27676440 | 27762957 | 74157.5 | retinoid X receptor alpha |  |  |  |  |  |
| rs4223708 | rs3694208 | -0.007 | 0.19421649 | -0.013 | 0.09656639 | 0.061 | 0.00243123 | 8.561 | 3 | 14.696 | 8 | Stmn2 | 8509360 | 8561606 | 25495 | stathmin-like 2 |  |  |  |  |  |
| rs4223708 | rs4140004 | -0.003 | 0.58916257 | -0.014 | 0.14059881 | 0.071 | 0.00939895 | 8.561 | 3 | 22.533 | 8 | Stmn2 | 8509360 | 8561606 | 25495 | stathmin-like 2 |  |  |  |  |  |
| rs3689073 | rs13479753 | -0.002 | 0.70929773 | 0.188 | 1.78E-10 | -0.149 | 0.00223861 | 7.385 | 3 | 50.552 | 8 | Pkia | 7366669 | 7445366 | 20762.5 | protein kinase inhibitor, alpha |  |  |  |  |  |
| rs13476961 | rs6322205 | -0.002 | 0.70003469 | 0.188 | 1.80E-10 | -0.149 | 0.00227424 | 6.808 | 3 | 51.103 | 8 | Olfr289-ps1 | 6751704 | 6752072 | 56067 |  |  |  |  |  |  |
| rs3725706 | rs13480205 | -0.045 | 1.86E-08 | -0.038 | 3.78E-09 | 0.043 | 0.00105522 | 41.747 | 3 | 54.530 | 9 | D3Ertd751e | 41742611 | 41803320 | 26432.5 | DNA segment, Chr 3, ERATO Doi 751, expressed |  |  |  |  |  |
| rs6324747 | rs13480302 | -0.035 | 3.65E-08 | -0.038 | 5.36E-07 | 0.043 | 0.00222863 | 40.032 | 3 | 80.733 | 9 | Gm42785 | 39823577 | 39827411 | 206985 |  |  |  |  |  |  |
| rs3672384 | rs3670195 | 0.024 | 1.06E-05 | 0.043 | 1.52E-09 | -0.049 | 0.00092982 | 62.881 | 3 | 81.266 | 9 | Gm9701 | 62923807 | 62924233 | 43313 |  |  |  |  |  |  |
| rs6175921 | rs13480302 | -0.032 | 1.08E-07 | -0.037 | 6.49E-07 | 0.042 | 0.00175799 | 67.371 | 4 | 80.733 | 9 | Hmgb1-rs18 | 67518306 | 67518932 | 148012 |  |  |  |  |  |  |
| rs3718146 | rs13480302 | -0.032 | 1.04E-07 | -0.037 | 6.48E-07 | 0.042 | 0.00170177 | 67.229 | 4 | 80.733 | 9 | Gm11403 | 67165479 | 67165847 | 63790 |  |  |  |  |  |  |
| rs13477755 | rs13480302 | -0.032 | 1.20E-07 | -0.037 | 6.45E-07 | 0.042 | 0.00160296 | 67.912 | 4 | 80.733 | 9 | Gm11249 | 67930881 | 67931739 | 18963 |  |  |  |  |  |  |
| rs13478570 | rs13480638 | 0.033 | 2.12E-06 | 0.019 | 0.00674874 | -0.039 | 0.00273646 | 142.647 | 5 | 69.445 | 10 | Wipi2 | 142627698 | 142670588 | 1882 | WD repeat domain, phosphoinositide interacting 2 |  |  |  |  |  |
| rs3686975 | rs3708535 | -0.017 | 0.00448946 | -0.026 | 0.00023963 | 0.043 | 4.50E-05 | 109.980 | 7 | 86.839 | 14 | Tmem41b | 109972187 | 109986929 | 187 | transmembrane protein 41B |  |  |  |  |  |
| rs13479196 | rs13479566 | 0.030 | 1.53E-06 | 0.033 | 1.67E-06 | -0.051 | 1.65E-05 | 37.481 | 7 | 142.463 | 7 | Gm44806 | 37465120 | 37467422 | 14526 |  |  |  |  |  |  |
| rs13479757 | rs3679439 | 0.136 | 2.93E-09 | -0.004 | 0.56324156 | -0.140 | 0.00657629 | 51.455 | 8 | 97.129 | 8 | Gm17911 | 51453395 | 51454271 | 1071 |  |  |  |  |  |  |
| rs3697959 | rs3725510 | 0.129 | 5.13E-09 | 0.004 | 0.56694653 | -0.143 | 0.00164256 | 50.837 | 8 | 102.238 | 8 | C130073E24Rik | 50743291 | 50916997 | 6947 |  |  |  |  |  |  |
| rs6322205 | rs3664869 | 0.136 | 2.93E-09 | -0.004 | 0.56324156 | -0.140 | 0.00657629 | 51.103 | 8 | 96.886 | 8 | Gm45386 | 51137630 | 51137847 | 34585.5 |  |  |  |  |  |  |
| rs13479757 | rs3664869 | 0.136 | 2.93E-09 | -0.004 | 0.56324156 | -0.140 | 0.00657629 | 51.455 | 8 | 96.886 | 8 | Gm17911 | 51453395 | 51454271 | 1071 |  |  |  |  |  |  |
| rs13479755 | rs3663506 | 0.138 | 1.77E-09 | 0.003 | 0.64241268 | -0.146 | 0.00087245 | 50.954 | 8 | 97.709 | 8 | C130073E24Rik | 50743291 | 50916997 | 124071 |  |  |  |  |  |  |
| rs6322205 | rs3663506 | 0.138 | 1.77E-09 | 0.003 | 0.64241268 | -0.146 | 0.00087245 | 51.103 | 8 | 97.709 | 8 | Gm45386 | 51137630 | 51137847 | 34585.5 |  |  |  |  |  |  |
| rs3697959 | rs3721390 | 0.129 | 5.13E-09 | 0.004 | 0.56694653 | -0.143 | 0.00164256 | 50.837 | 8 | 100.433 | 8 | C130073E24Rik | 50743291 | 50916997 | 6947 |  |  |  |  |  |  |
| rs6244635 | rs3725510 | 0.129 | 5.13E-09 | 0.004 | 0.56694653 | -0.143 | 0.00164256 | 51.213 | 8 | 102.238 | 8 | Gm32975 | 51177683 | 51180608 | 34251.5 |  |  |  |  |  |  |
| rs13479654 | rs13479993 | 0.053 | 4.37E-06 | 0.000 | 0.98580609 | -0.124 | 0.00654386 | 25.178 | 8 | 112.929 | 8 | Mir8108 | 25159125 | 25159233 | 18936 |  |  |  |  |  |  |
| rs3697959 | rs13479993 | 0.124 | 3.89E-09 | -0.003 | 0.60336264 | -0.160 | 0.00047341 | 50.837 | 8 | 112.929 | 8 | C130073E24Rik | 50743291 | 50916997 |  |  |  |  |  |  |  |
| rs13480793 | rs13482148 | 0.039 | 3.05E-08 | 0.023 | 0.00013031 | -0.044 | 0.00299396 | 121.028 | 10 | 41.187 | 14 | Gm48341 | 120984756 | 121001061 |  |  |  |  |  |  |  |
| rs13481249 | rs3699140 | -0.029 | 6.26E-05 | -0.020 | 0.0012578 | 0.046 | 0.00560757 | 116.542 | 11 | 84.871 | 14 | Sphk1 | 116530925 | 116536675 |  |  |  |  |  |  |  |
| rs3670719 | rs13482019 | 0.056 | 0.00019547 | -0.006 | 0.19659078 | -0.082 | 0.00091106 | 69.092 | 13 | 112.768 | 13 | Gm26844 | 69141229 | 69143965 |  |  |  |  |  |  |  |
| rs3670719 | rs13482018 | 0.055 | 0.00028447 | -0.006 | 0.1930375 | -0.080 | 0.00257419 | 69.092 | 13 | 112.636 | 13 | Gm26844 | 69141229 | 69143965 |  |  |  |  |  |  |  |
| rs6367778 | rs3719853 | 0.051 | 0.00021723 | 0.003 | 0.43384516 | -0.081 | 0.00017755 | 69.367 | 13 | 113.197 | 13 | Gm4812 | 69345655 | 69346647 |  |  |  |  |  |  |  |
| rs3670719 | rs13482016 | -0.100 | 3.28E-13 | 0.007 | 0.08895526 | 0.078 | 0.00385739 | 69.092 | 13 | 111.953 | 13 | Gm26844 | 69141229 | 69143965 |  |  |  |  |  |  |  |
| rs3670719 | rs3719853 | 0.063 | 3.37E-05 | 0.004 | 0.40127032 | -0.090 | 8.28E-06 | 69.092 | 13 | 113.197 | 13 | Gm26844 | 69141229 | 69143965 |  |  |  |  |  |  |  |
| rs3670719 | rs4230072 | -0.099 | 5.22E-13 | 0.008 | 0.05367099 | 0.077 | 0.00598304 | 69.092 | 13 | 112.738 | 13 | Gm26844 | 69141229 | 69143965 |  |  |  |  |  |  |  |
| rs6367778 | rs13483571 | -0.015 | 0.07463469 | 0.022 | 0.02456875 | -0.213 | 0.00327398 | 69.367 | 13 | 24.352 | 19 | Gm4812 | 69345655 | 69346647 |  |  |  |  |  |  |  |
