## Supplementary Table S3 for "Detecting epistatic interactions in genomic data using Random Forests"

**Supplementary Table S3.** List of interacting murine SNP for the W6W trait and associated genomic features

| **Input** | **MGI Gene/Marker ID** | **Symbol** | **Name** | **Feature Type** | **Ensembl ID** | **Entrez Gene ID** | **GO ID** | **Term** |
| --- | --- | --- | --- | --- | --- | --- | --- | --- |
| rs3670719 | No associated gene |  |  |  |  |  |  |  |
| rs13476400 | MGI:5676472 | Cpgi11386 | CpG island 11386 | CpG island |  |  |  |  |
| rs13476400 | MGI:2153089 | Mrps2 | mitochondrial ribosomal protein S2 | protein coding gene | ENSMUSG00000035772 | 118451 | GO:0005622 | intracellular |
| rs13476400 | MGI:2153089 | Mrps2 | mitochondrial ribosomal protein S2 | protein coding gene | ENSMUSG00000035772 | 118451 | GO:0061668 | mitochondrial ribosome assembly |
| rs13476400 | MGI:2153089 | Mrps2 | mitochondrial ribosomal protein S2 | protein coding gene | ENSMUSG00000035772 | 118451 | GO:0005763 | mitochondrial small ribosomal subunit |
| rs13476400 | MGI:2153089 | Mrps2 | mitochondrial ribosomal protein S2 | protein coding gene | ENSMUSG00000035772 | 118451 | GO:0005763 | mitochondrial small ribosomal subunit |
| rs13476400 | MGI:2153089 | Mrps2 | mitochondrial ribosomal protein S2 | protein coding gene | ENSMUSG00000035772 | 118451 | GO:0032543 | mitochondrial translation |
| rs13476400 | MGI:2153089 | Mrps2 | mitochondrial ribosomal protein S2 | protein coding gene | ENSMUSG00000035772 | 118451 | GO:0005739 | mitochondrion |
| rs13476400 | MGI:2153089 | Mrps2 | mitochondrial ribosomal protein S2 | protein coding gene | ENSMUSG00000035772 | 118451 | GO:0005739 | mitochondrion |
| rs13476400 | MGI:2153089 | Mrps2 | mitochondrial ribosomal protein S2 | protein coding gene | ENSMUSG00000035772 | 118451 | GO:0005840 | ribosome |
| rs13476400 | MGI:2153089 | Mrps2 | mitochondrial ribosomal protein S2 | protein coding gene | ENSMUSG00000035772 | 118451 | GO:0015935 | small ribosomal subunit |
| rs13476400 | MGI:2153089 | Mrps2 | mitochondrial ribosomal protein S2 | protein coding gene | ENSMUSG00000035772 | 118451 | GO:0003735 | structural constituent of ribosome |
| rs13476400 | MGI:2153089 | Mrps2 | mitochondrial ribosomal protein S2 | protein coding gene | ENSMUSG00000035772 | 118451 | GO:0006412 | translation |
| rs13476400 | MGI:5944695 | Tssr16928 | transcription start site region 16928 | TSS region |  |  |  |  |
| rs13476400 | MGI:5944696 | Tssr16929 | transcription start site region 16929 | TSS region |  |  |  |  |
| rs13476400 | MGI:5944697 | Tssr16930 | transcription start site region 16930 | TSS region |  |  |  |  |
| rs13476400 | MGI:5944698 | Tssr16931 | transcription start site region 16931 | TSS region |  |  |  |  |
| rs13476402 | MGI:2682639 | Ddx31 | DEAD/H box helicase 31 | protein coding gene | ENSMUSG00000026806 | 227674 | GO:0005524 | ATP binding |
| rs13476402 | MGI:2682639 | Ddx31 | DEAD/H box helicase 31 | protein coding gene | ENSMUSG00000026806 | 227674 | GO:0005794 | Golgi apparatus |
| rs13476402 | MGI:2682639 | Ddx31 | DEAD/H box helicase 31 | protein coding gene | ENSMUSG00000026806 | 227674 | GO:0004386 | helicase activity |
| rs13476402 | MGI:2682639 | Ddx31 | DEAD/H box helicase 31 | protein coding gene | ENSMUSG00000026806 | 227674 | GO:0016787 | hydrolase activity |
| rs13476402 | MGI:2682639 | Ddx31 | DEAD/H box helicase 31 | protein coding gene | ENSMUSG00000026806 | 227674 | GO:0043231 | intracellular membrane-bounded organelle |
| rs13476402 | MGI:2682639 | Ddx31 | DEAD/H box helicase 31 | protein coding gene | ENSMUSG00000026806 | 227674 | GO:0003676 | nucleic acid binding |
| rs13476402 | MGI:2682639 | Ddx31 | DEAD/H box helicase 31 | protein coding gene | ENSMUSG00000026806 | 227674 | GO:0005730 | nucleolus |
| rs13476402 | MGI:2682639 | Ddx31 | DEAD/H box helicase 31 | protein coding gene | ENSMUSG00000026806 | 227674 | GO:0000166 | nucleotide binding |
| rs13476402 | MGI:2682639 | Ddx31 | DEAD/H box helicase 31 | protein coding gene | ENSMUSG00000026806 | 227674 | GO:0042254 | ribosome biogenesis |
| rs13476402 | MGI:2682639 | Ddx31 | DEAD/H box helicase 31 | protein coding gene | ENSMUSG00000026806 | 227674 | GO:0003723 | RNA binding |
| rs6367778 | No associated gene | |  |  |  |  |  |  |
| rs3023797 | MGI:3649783 | Gm13393 | predicted gene 13393 | lncRNA gene | ENSMUSG00000086519 | |  |  |
| rs3023797 | MGI:105044 | Ttf1 | transcription termination factor, RNA polymerase I | protein coding gene | ENSMUSG00000026803 | 22130 | GO:0003682 | chromatin binding |
| rs3023797 | MGI:105044 | Ttf1 | transcription termination factor, RNA polymerase I | protein coding gene | ENSMUSG00000026803 | 22130 | GO:0006338 | chromatin remodeling |
| rs3023797 | MGI:105044 | Ttf1 | transcription termination factor, RNA polymerase I | protein coding gene | ENSMUSG00000026803 | 22130 | GO:0003677 | DNA binding |
| rs3023797 | MGI:105044 | Ttf1 | transcription termination factor, RNA polymerase I | protein coding gene | ENSMUSG00000026803 | 22130 | GO:0006353 | DNA-templated transcription, termination |
| rs3023797 | MGI:105044 | Ttf1 | transcription termination factor, RNA polymerase I | protein coding gene | ENSMUSG00000026803 | 22130 | GO:0008156 | negative regulation of DNA replication |
| rs3023797 | MGI:105044 | Ttf1 | transcription termination factor, RNA polymerase I | protein coding gene | ENSMUSG00000026803 | 22130 | GO:0005634 | nucleus |
| rs3023797 | MGI:105044 | Ttf1 | transcription termination factor, RNA polymerase I | protein coding gene | ENSMUSG00000026803 | 22130 | GO:0005515 | protein binding |
| rs3023797 | MGI:105044 | Ttf1 | transcription termination factor, RNA polymerase I | protein coding gene | ENSMUSG00000026803 | 22130 | GO:0006363 | termination of RNA polymerase I transcription |
| rs3023797 | MGI:105044 | Ttf1 | transcription termination factor, RNA polymerase I | protein coding gene | ENSMUSG00000026803 | 22130 | GO:0006361 | transcription initiation from RNA polymerase I promoter |
| rs4223708 | MGI:98241 | Stmn2 | stathmin-like 2 | protein coding gene | ENSMUSG00000027500 | 20257 | GO:0048306 | calcium-dependent protein binding |
| rs4223708 | MGI:98241 | Stmn2 | stathmin-like 2 | protein coding gene | ENSMUSG00000027500 | 20257 | GO:0042995 | cell projection |
| rs4223708 | MGI:98241 | Stmn2 | stathmin-like 2 | protein coding gene | ENSMUSG00000027500 | 20257 | GO:1990090 | cellular response to nerve growth factor stimulus |
| rs4223708 | MGI:98241 | Stmn2 | stathmin-like 2 | protein coding gene | ENSMUSG00000027500 | 20257 | GO:0005737 | cytoplasm |
| rs4223708 | MGI:98241 | Stmn2 | stathmin-like 2 | protein coding gene | ENSMUSG00000027500 | 20257 | GO:0005768 | endosome |
| rs4223708 | MGI:98241 | Stmn2 | stathmin-like 2 | protein coding gene | ENSMUSG00000027500 | 20257 | GO:0005794 | Golgi apparatus |
| rs4223708 | MGI:98241 | Stmn2 | stathmin-like 2 | protein coding gene | ENSMUSG00000027500 | 20257 | GO:0030426 | growth cone |
| rs4223708 | MGI:98241 | Stmn2 | stathmin-like 2 | protein coding gene | ENSMUSG00000027500 | 20257 | GO:0030426 | growth cone |
| rs4223708 | MGI:98241 | Stmn2 | stathmin-like 2 | protein coding gene | ENSMUSG00000027500 | 20257 | GO:0030027 | lamellipodium |
| rs4223708 | MGI:98241 | Stmn2 | stathmin-like 2 | protein coding gene | ENSMUSG00000027500 | 20257 | GO:0016020 | membrane |
| rs4223708 | MGI:98241 | Stmn2 | stathmin-like 2 | protein coding gene | ENSMUSG00000027500 | 20257 | GO:0007026 | negative regulation of microtubule depolymerization |
| rs4223708 | MGI:98241 | Stmn2 | stathmin-like 2 | protein coding gene | ENSMUSG00000027500 | 20257 | GO:0031115 | negative regulation of microtubule polymerization |
| rs4223708 | MGI:98241 | Stmn2 | stathmin-like 2 | protein coding gene | ENSMUSG00000027500 | 20257 | GO:0010977 | negative regulation of neuron projection development |
| rs4223708 | MGI:98241 | Stmn2 | stathmin-like 2 | protein coding gene | ENSMUSG00000027500 | 20257 | GO:0043025 | neuronal cell body |
| rs4223708 | MGI:98241 | Stmn2 | stathmin-like 2 | protein coding gene | ENSMUSG00000027500 | 20257 | GO:0043005 | neuron projection |
| rs4223708 | MGI:98241 | Stmn2 | stathmin-like 2 | protein coding gene | ENSMUSG00000027500 | 20257 | GO:0048471 | perinuclear region of cytoplasm |
| rs4223708 | MGI:98241 | Stmn2 | stathmin-like 2 | protein coding gene | ENSMUSG00000027500 | 20257 | GO:0031117 | positive regulation of microtubule depolymerization |
| rs4223708 | MGI:98241 | Stmn2 | stathmin-like 2 | protein coding gene | ENSMUSG00000027500 | 20257 | GO:0010976 | positive regulation of neuron projection development |
| rs4223708 | MGI:98241 | Stmn2 | stathmin-like 2 | protein coding gene | ENSMUSG00000027500 | 20257 | GO:0005515 | protein binding |
| rs4223708 | MGI:98241 | Stmn2 | stathmin-like 2 | protein coding gene | ENSMUSG00000027500 | 20257 | GO:0031982 | vesicle |
| rs4223708 | MGI:5956097 | Tssr28330 | transcription start site region 28330 | TSS region |  |  |  |  |
| rs4223708 | MGI:5956098 | Tssr28331 | transcription start site region 28331 | TSS region |  |  |  |  |
| rs13479757 | MGI:5010096 | Gm17911 | predicted gene, 17911 | pseudogene | ENSMUSG00000110085 | 100416081 |  |  |
| rs6308258 | MGI:107485 | Ralgds | ral guanine nucleotide dissociation stimulator | protein coding gene | ENSMUSG00000026821 | 19730 | GO:0005903 | brush border |
| rs6308258 | MGI:107485 | Ralgds | ral guanine nucleotide dissociation stimulator | protein coding gene | ENSMUSG00000026821 | 19730 | GO:0005737 | cytoplasm |
| rs6308258 | MGI:107485 | Ralgds | ral guanine nucleotide dissociation stimulator | protein coding gene | ENSMUSG00000026821 | 19730 | GO:0005829 | cytosol |
| rs6308258 | MGI:107485 | Ralgds | ral guanine nucleotide dissociation stimulator | protein coding gene | ENSMUSG00000026821 | 19730 | GO:0030695 | GTPase regulator activity |
| rs6308258 | MGI:107485 | Ralgds | ral guanine nucleotide dissociation stimulator | protein coding gene | ENSMUSG00000026821 | 19730 | GO:0005085 | guanyl-nucleotide exchange factor activity |
| rs6308258 | MGI:107485 | Ralgds | ral guanine nucleotide dissociation stimulator | protein coding gene | ENSMUSG00000026821 | 19730 | GO:0005622 | intracellular |
| rs6308258 | MGI:107485 | Ralgds | ral guanine nucleotide dissociation stimulator | protein coding gene | ENSMUSG00000026821 | 19730 | GO:0005634 | nucleus |
| rs6308258 | MGI:107485 | Ralgds | ral guanine nucleotide dissociation stimulator | protein coding gene | ENSMUSG00000026821 | 19730 | GO:0005515 | protein binding |
| rs6308258 | MGI:107485 | Ralgds | ral guanine nucleotide dissociation stimulator | protein coding gene | ENSMUSG00000026821 | 19730 | GO:0007165 | signal transduction |
| rs6308258 | MGI:107485 | Ralgds | ral guanine nucleotide dissociation stimulator | protein coding gene | ENSMUSG00000026821 | 19730 | GO:0007264 | small GTPase mediated signal transduction |
| rs6308258 | MGI:5944718 | Tssr16951 | transcription start site region 16951 | TSS region |  |  |  |  |
| rs3686975 | MGI:1289225 | Tmem41b | transmembrane protein 41B | protein coding gene | ENSMUSG00000047554 | 233724 | GO:0000045 | autophagosome assembly |
| rs3686975 | MGI:1289225 | Tmem41b | transmembrane protein 41B | protein coding gene | ENSMUSG00000047554 | 233724 | GO:0006914 | autophagy |
| rs3686975 | MGI:1289225 | Tmem41b | transmembrane protein 41B | protein coding gene | ENSMUSG00000047554 | 233724 | GO:0005783 | endoplasmic reticulum |
| rs3686975 | MGI:1289225 | Tmem41b | transmembrane protein 41B | protein coding gene | ENSMUSG00000047554 | 233724 | GO:0005789 | endoplasmic reticulum membrane |
| rs3686975 | MGI:1289225 | Tmem41b | transmembrane protein 41B | protein coding gene | ENSMUSG00000047554 | 233724 | GO:0016021 | integral component of membrane |
| rs3686975 | MGI:1289225 | Tmem41b | transmembrane protein 41B | protein coding gene | ENSMUSG00000047554 | 233724 | GO:0016020 | membrane |
| rs3686975 | MGI:1289225 | Tmem41b | transmembrane protein 41B | protein coding gene | ENSMUSG00000047554 | 233724 | GO:0044233 | mitochondria-associated endoplasmic reticulum membrane |
| rs3686975 | MGI:1289225 | Tmem41b | transmembrane protein 41B | protein coding gene | ENSMUSG00000047554 | 233724 | GO:0044233 | mitochondria-associated endoplasmic reticulum membrane |
| rs3686975 | MGI:1289225 | Tmem41b | transmembrane protein 41B | protein coding gene | ENSMUSG00000047554 | 233724 | GO:0003674 | molecular_function |
| rs3686975 | MGI:1289225 | Tmem41b | transmembrane protein 41B | protein coding gene | ENSMUSG00000047554 | 233724 | GO:0007399 | nervous system development |
| rs3697959 | MGI:1922799 | 1700019L22Rik | RIKEN cDNA 1700019L22 gene | lncRNA gene | | 75549 |  |  |
| rs3697959 | MGI:2442759 | C130073E24Rik | RIKEN cDNA C130073E24 gene | lncRNA gene | ENSMUSG00000110246 | 319805 |  |  |
| rs3697959 | MGI:3764281 | D8Dcr17 | DNA Segment, Chr 8, Derry C. Roopenian 17 | DNA segment | |  |  |  |
| rs3697959 | MGI:100205 | D8Mit205 | DNA segment, Chr 8, Massachusetts Institute of Technology 205 | DNA segment | | 61647 |  |  |
| rs6175921 | No associated gene | |  |  |  |  |  |  |
| rs13478570 | MGI:1923831 | Wipi2 | WD repeat domain, phosphoinositide interacting 2 | protein coding gene | ENSMUSG00000029578 | 74781 | GO:0005776 | autophagosome |
| rs13478570 | MGI:1923831 | Wipi2 | WD repeat domain, phosphoinositide interacting 2 | protein coding gene | ENSMUSG00000029578 | 74781 | GO:0000045 | autophagosome assembly |
| rs13478570 | MGI:1923831 | Wipi2 | WD repeat domain, phosphoinositide interacting 2 | protein coding gene | ENSMUSG00000029578 | 74781 | GO:0006914 | autophagy |
| rs13478570 | MGI:1923831 | Wipi2 | WD repeat domain, phosphoinositide interacting 2 | protein coding gene | ENSMUSG00000029578 | 74781 | GO:0009267 | cellular response to starvation |
| rs13478570 | MGI:1923831 | Wipi2 | WD repeat domain, phosphoinositide interacting 2 | protein coding gene | ENSMUSG00000029578 | 74781 | GO:0005829 | cytosol |
| rs13478570 | MGI:1923831 | Wipi2 | WD repeat domain, phosphoinositide interacting 2 | protein coding gene | ENSMUSG00000029578 | 74781 | GO:0008289 | lipid binding |
| rs13478570 | MGI:1923831 | Wipi2 | WD repeat domain, phosphoinositide interacting 2 | protein coding gene | ENSMUSG00000029578 | 74781 | GO:0016020 | membrane |
| rs13478570 | MGI:1923831 | Wipi2 | WD repeat domain, phosphoinositide interacting 2 | protein coding gene | ENSMUSG00000029578 | 74781 | GO:0005654 | nucleoplasm |
| rs13478570 | MGI:1923831 | Wipi2 | WD repeat domain, phosphoinositide interacting 2 | protein coding gene | ENSMUSG00000029578 | 74781 | GO:0000407 | phagophore assembly site |
| rs13478570 | MGI:1923831 | Wipi2 | WD repeat domain, phosphoinositide interacting 2 | protein coding gene | ENSMUSG00000029578 | 74781 | GO:0000407 | phagophore assembly site |
| rs13478570 | MGI:1923831 | Wipi2 | WD repeat domain, phosphoinositide interacting 2 | protein coding gene | ENSMUSG00000029578 | 74781 | GO:0034045 | phagophore assembly site membrane |
| rs13478570 | MGI:1923831 | Wipi2 | WD repeat domain, phosphoinositide interacting 2 | protein coding gene | ENSMUSG00000029578 | 74781 | GO:0080025 | phosphatidylinositol-3,5-bisphosphate binding |
| rs13478570 | MGI:1923831 | Wipi2 | WD repeat domain, phosphoinositide interacting 2 | protein coding gene | ENSMUSG00000029578 | 74781 | GO:0032266 | phosphatidylinositol-3-phosphate binding |
| rs13478570 | MGI:1923831 | Wipi2 | WD repeat domain, phosphoinositide interacting 2 | protein coding gene | ENSMUSG00000029578 | 74781 | GO:0032266 | phosphatidylinositol-3-phosphate binding |
| rs13478570 | MGI:1923831 | Wipi2 | WD repeat domain, phosphoinositide interacting 2 | protein coding gene | ENSMUSG00000029578 | 74781 | GO:0010314 | phosphatidylinositol-5-phosphate binding |
| rs13478570 | MGI:1923831 | Wipi2 | WD repeat domain, phosphoinositide interacting 2 | protein coding gene | ENSMUSG00000029578 | 74781 | GO:0005515 | protein binding |
| rs13478570 | MGI:1923831 | Wipi2 | WD repeat domain, phosphoinositide interacting 2 | protein coding gene | ENSMUSG00000029578 | 74781 | GO:0032991 | protein-containing complex |
| rs13478570 | MGI:1923831 | Wipi2 | WD repeat domain, phosphoinositide interacting 2 | protein coding gene | ENSMUSG00000029578 | 74781 | GO:0061739 | protein lipidation involved in autophagosome assembly |
| rs13478570 | MGI:1923831 | Wipi2 | WD repeat domain, phosphoinositide interacting 2 | protein coding gene | ENSMUSG00000029578 | 74781 | GO:0034497 | protein localization to phagophore assembly site |
| rs13478570 | MGI:1923831 | Wipi2 | WD repeat domain, phosphoinositide interacting 2 | protein coding gene | ENSMUSG00000029578 | 74781 | GO:0034497 | protein localization to phagophore assembly site |
| rs13478570 | MGI:1923831 | Wipi2 | WD repeat domain, phosphoinositide interacting 2 | protein coding gene | ENSMUSG00000029578 | 74781 | GO:0098792 | xenophagy |
| rs13480793 | MGI:5623669 | Gm40784 | predicted gene, 40784 | lncRNA gene | | 105245309 |  |  |
| rs13481249 | MGI:6039706 | Tssr111939 | transcription start site region 111939 | TSS region |  |  |  |  |
| rs13481249 | MGI:2444266 | Ube2o | ubiquitin-conjugating enzyme E2O | protein coding gene | ENSMUSG00000020802 | 217342 | GO:0005524 | ATP binding |
| rs13481249 | MGI:2444266 | Ube2o | ubiquitin-conjugating enzyme E2O | protein coding gene | ENSMUSG00000020802 | 217342 | GO:0005737 | cytoplasm |
| rs13481249 | MGI:2444266 | Ube2o | ubiquitin-conjugating enzyme E2O | protein coding gene | ENSMUSG00000020802 | 217342 | GO:0016604 | nuclear body |
| rs13481249 | MGI:2444266 | Ube2o | ubiquitin-conjugating enzyme E2O | protein coding gene | ENSMUSG00000020802 | 217342 | GO:0005654 | nucleoplasm |
| rs13481249 | MGI:2444266 | Ube2o | ubiquitin-conjugating enzyme E2O | protein coding gene | ENSMUSG00000020802 | 217342 | GO:0000166 | nucleotide binding |
| rs13481249 | MGI:2444266 | Ube2o | ubiquitin-conjugating enzyme E2O | protein coding gene | ENSMUSG00000020802 | 217342 | GO:0005634 | nucleus |
| rs13481249 | MGI:2444266 | Ube2o | ubiquitin-conjugating enzyme E2O | protein coding gene | ENSMUSG00000020802 | 217342 | GO:0030513 | positive regulation of BMP signaling pathway |
| rs13481249 | MGI:2444266 | Ube2o | ubiquitin-conjugating enzyme E2O | protein coding gene | ENSMUSG00000020802 | 217342 | GO:0005515 | protein binding |
| rs13481249 | MGI:2444266 | Ube2o | ubiquitin-conjugating enzyme E2O | protein coding gene | ENSMUSG00000020802 | 217342 | GO:0070534 | protein K63-linked ubiquitination |
| rs13481249 | MGI:2444266 | Ube2o | ubiquitin-conjugating enzyme E2O | protein coding gene | ENSMUSG00000020802 | 217342 | GO:0006513 | protein monoubiquitination |
| rs13481249 | MGI:2444266 | Ube2o | ubiquitin-conjugating enzyme E2O | protein coding gene | ENSMUSG00000020802 | 217342 | GO:0042147 | retrograde transport, endosome to Golgi |
| rs13481249 | MGI:2444266 | Ube2o | ubiquitin-conjugating enzyme E2O | protein coding gene | ENSMUSG00000020802 | 217342 | GO:0016740 | transferase activity |
| rs13481249 | MGI:2444266 | Ube2o | ubiquitin-conjugating enzyme E2O | protein coding gene | ENSMUSG00000020802 | 217342 | GO:0061631 | ubiquitin conjugating enzyme activity |
| rs13481249 | MGI:2444266 | Ube2o | ubiquitin-conjugating enzyme E2O | protein coding gene | ENSMUSG00000020802 | 217342 | GO:0061630 | ubiquitin protein ligase activity |
| rs13481249 | MGI:2444266 | Ube2o | ubiquitin-conjugating enzyme E2O | protein coding gene | ENSMUSG00000020802 | 217342 | GO:0004842 | ubiquitin-protein transferase activity |
| rs13469412 | MGI:1913420 | Chchd5 | coiled-coil-helix-coiled-coil-helix domain containing 5 | protein coding gene | ENSMUSG00000037938 | 66170 | GO:0005739 | mitochondrion |
| rs13469412 | MGI:1913420 | Chchd5 | coiled-coil-helix-coiled-coil-helix domain containing 5 | protein coding gene | ENSMUSG00000037938 | 66170 | GO:0003674 | molecular_function |
| rs13479196 | MGI:5993696 | Tssr65929 | transcription start site region 65929 | TSS region |  |  |  |  |
| rs13479196 | MGI:5999835 | Tssr72068 | transcription start site region 72068 | TSS region |  |  |  |  |
| rs13479196 | MGI:1926102 | Zfp536 | zinc finger protein 536 | protein coding gene | ENSMUSG00000043456 | 243937 | GO:0003677 | DNA binding |
| rs13479196 | MGI:1926102 | Zfp536 | zinc finger protein 536 | protein coding gene | ENSMUSG00000043456 | 243937 | GO:0001227 | DNA-binding transcription repressor activity, RNA polymerase II-specific |
| rs13479196 | MGI:1926102 | Zfp536 | zinc finger protein 536 | protein coding gene | ENSMUSG00000043456 | 243937 | GO:0046872 | metal ion binding |
| rs13479196 | MGI:1926102 | Zfp536 | zinc finger protein 536 | protein coding gene | ENSMUSG00000043456 | 243937 | GO:0045665 | negative regulation of neuron differentiation |
| rs13479196 | MGI:1926102 | Zfp536 | zinc finger protein 536 | protein coding gene | ENSMUSG00000043456 | 243937 | GO:0048387 | negative regulation of retinoic acid receptor signaling pathway |
| rs13479196 | MGI:1926102 | Zfp536 | zinc finger protein 536 | protein coding gene | ENSMUSG00000043456 | 243937 | GO:0000122 | negative regulation of transcription by RNA polymerase II |
| rs13479196 | MGI:1926102 | Zfp536 | zinc finger protein 536 | protein coding gene | ENSMUSG00000043456 | 243937 | GO:0003676 | nucleic acid binding |
| rs13479196 | MGI:1926102 | Zfp536 | zinc finger protein 536 | protein coding gene | ENSMUSG00000043456 | 243937 | GO:0005634 | nucleus |
| rs13479196 | MGI:1926102 | Zfp536 | zinc finger protein 536 | protein coding gene | ENSMUSG00000043456 | 243937 | GO:0044323 | retinoic acid-responsive element binding |
| rs13479196 | MGI:1926102 | Zfp536 | zinc finger protein 536 | protein coding gene | ENSMUSG00000043456 | 243937 | GO:0000978 | RNA polymerase II cis-regulatory region sequence-specific DNA binding |
| rs3163007 | MGI:1935151 | Klf7 | Kruppel-like factor 7 (ubiquitous) | protein coding gene | ENSMUSG00000025959 | 93691 | GO:0007411 | axon guidance |
| rs3163007 | MGI:1935151 | Klf7 | Kruppel-like factor 7 (ubiquitous) | protein coding gene | ENSMUSG00000025959 | 93691 | GO:0007409 | axonogenesis |
| rs3163007 | MGI:1935151 | Klf7 | Kruppel-like factor 7 (ubiquitous) | protein coding gene | ENSMUSG00000025959 | 93691 | GO:0048813 | dendrite morphogenesis |
| rs3163007 | MGI:1935151 | Klf7 | Kruppel-like factor 7 (ubiquitous) | protein coding gene | ENSMUSG00000025959 | 93691 | GO:0003677 | DNA binding |
| rs3163007 | MGI:1935151 | Klf7 | Kruppel-like factor 7 (ubiquitous) | protein coding gene | ENSMUSG00000025959 | 93691 | GO:0003677 | DNA binding |
| rs3163007 | MGI:1935151 | Klf7 | Kruppel-like factor 7 (ubiquitous) | protein coding gene | ENSMUSG00000025959 | 93691 | GO:0001228 | DNA-binding transcription activator activity, RNA polymerase II-specific |
| rs3163007 | MGI:1935151 | Klf7 | Kruppel-like factor 7 (ubiquitous) | protein coding gene | ENSMUSG00000025959 | 93691 | GO:0003700 | DNA-binding transcription factor activity |
| rs3163007 | MGI:1935151 | Klf7 | Kruppel-like factor 7 (ubiquitous) | protein coding gene | ENSMUSG00000025959 | 93691 | GO:0000981 | DNA-binding transcription factor activity, RNA polymerase II-specific |
| rs3163007 | MGI:1935151 | Klf7 | Kruppel-like factor 7 (ubiquitous) | protein coding gene | ENSMUSG00000025959 | 93691 | GO:0042593 | glucose homeostasis |
| rs3163007 | MGI:1935151 | Klf7 | Kruppel-like factor 7 (ubiquitous) | protein coding gene | ENSMUSG00000025959 | 93691 | GO:0046872 | metal ion binding |
| rs3163007 | MGI:1935151 | Klf7 | Kruppel-like factor 7 (ubiquitous) | protein coding gene | ENSMUSG00000025959 | 93691 | GO:1904178 | negative regulation of adipose tissue development |
| rs3163007 | MGI:1935151 | Klf7 | Kruppel-like factor 7 (ubiquitous) | protein coding gene | ENSMUSG00000025959 | 93691 | GO:0061179 | negative regulation of insulin secretion involved in cellular response to glucose stimulus |
| rs3163007 | MGI:1935151 | Klf7 | Kruppel-like factor 7 (ubiquitous) | protein coding gene | ENSMUSG00000025959 | 93691 | GO:0000122 | negative regulation of transcription by RNA polymerase II |
| rs3163007 | MGI:1935151 | Klf7 | Kruppel-like factor 7 (ubiquitous) | protein coding gene | ENSMUSG00000025959 | 93691 | GO:0003676 | nucleic acid binding |
| rs3163007 | MGI:1935151 | Klf7 | Kruppel-like factor 7 (ubiquitous) | protein coding gene | ENSMUSG00000025959 | 93691 | GO:0005634 | nucleus |
| rs3163007 | MGI:1935151 | Klf7 | Kruppel-like factor 7 (ubiquitous) | protein coding gene | ENSMUSG00000025959 | 93691 | GO:0045944 | positive regulation of transcription by RNA polymerase II |
| rs3163007 | MGI:1935151 | Klf7 | Kruppel-like factor 7 (ubiquitous) | protein coding gene | ENSMUSG00000025959 | 93691 | GO:0045893 | positive regulation of transcription, DNA-templated |
| rs3163007 | MGI:1935151 | Klf7 | Kruppel-like factor 7 (ubiquitous) | protein coding gene | ENSMUSG00000025959 | 93691 | GO:0005515 | protein binding |
| rs3163007 | MGI:1935151 | Klf7 | Kruppel-like factor 7 (ubiquitous) | protein coding gene | ENSMUSG00000025959 | 93691 | GO:0045604 | regulation of epidermal cell differentiation |
| rs3163007 | MGI:1935151 | Klf7 | Kruppel-like factor 7 (ubiquitous) | protein coding gene | ENSMUSG00000025959 | 93691 | GO:0000978 | RNA polymerase II cis-regulatory region sequence-specific DNA binding |
| rs3163007 | MGI:5530766 | Mir6899 | microRNA 6899 | miRNA gene | ENSMUSG00000098487 | 102466760 |  |  |
| rs3163007 | MGI:5939670 | Tssr11903 | transcription start site region 11903 | TSS region |  |  |  |  |
| rs3162895 | MGI:5579687 | Gm28981 | predicted gene 28981 | lncRNA gene | ENSMUSG00000101895 | |  |  |
| rs3162895 | MGI:5595039 | Gm35880 | predicted gene, 35880 | lncRNA gene | | 102639609 |  |  |
| rs6212146 | No associated gene | |  |  |  |  |  |  |
| rs3689073 | MGI:104747 | Pkia | protein kinase inhibitor, alpha | protein coding gene | ENSMUSG00000027499 | 18767 | GO:0004862 | cAMP-dependent protein kinase inhibitor activity |
| rs3689073 | MGI:104747 | Pkia | protein kinase inhibitor, alpha | protein coding gene | ENSMUSG00000027499 | 18767 | GO:0004862 | cAMP-dependent protein kinase inhibitor activity |
| rs3689073 | MGI:104747 | Pkia | protein kinase inhibitor, alpha | protein coding gene | ENSMUSG00000027499 | 18767 | GO:0005737 | cytoplasm |
| rs3689073 | MGI:104747 | Pkia | protein kinase inhibitor, alpha | protein coding gene | ENSMUSG00000027499 | 18767 | GO:2000480 | negative regulation of cAMP-dependent protein kinase activity |
| rs3689073 | MGI:104747 | Pkia | protein kinase inhibitor, alpha | protein coding gene | ENSMUSG00000027499 | 18767 | GO:0043086 | negative regulation of catalytic activity |
| rs3689073 | MGI:104747 | Pkia | protein kinase inhibitor, alpha | protein coding gene | ENSMUSG00000027499 | 18767 | GO:0042308 | negative regulation of protein import into nucleus |
| rs3689073 | MGI:104747 | Pkia | protein kinase inhibitor, alpha | protein coding gene | ENSMUSG00000027499 | 18767 | GO:0006469 | negative regulation of protein kinase activity |
| rs3689073 | MGI:104747 | Pkia | protein kinase inhibitor, alpha | protein coding gene | ENSMUSG00000027499 | 18767 | GO:0000122 | negative regulation of transcription by RNA polymerase II |
| rs3689073 | MGI:104747 | Pkia | protein kinase inhibitor, alpha | protein coding gene | ENSMUSG00000027499 | 18767 | GO:0005634 | nucleus |
| rs3689073 | MGI:104747 | Pkia | protein kinase inhibitor, alpha | protein coding gene | ENSMUSG00000027499 | 18767 | GO:0005515 | protein binding |
| rs3689073 | MGI:104747 | Pkia | protein kinase inhibitor, alpha | protein coding gene | ENSMUSG00000027499 | 18767 | GO:0034236 | protein kinase A catalytic subunit binding |
| rs3689073 | MGI:104747 | Pkia | protein kinase inhibitor, alpha | protein coding gene | ENSMUSG00000027499 | 18767 | GO:0034236 | protein kinase A catalytic subunit binding |
| rs3689073 | MGI:104747 | Pkia | protein kinase inhibitor, alpha | protein coding gene | ENSMUSG00000027499 | 18767 | GO:0004860 | protein kinase inhibitor activity |
| rs3689073 | MGI:104747 | Pkia | protein kinase inhibitor, alpha | protein coding gene | ENSMUSG00000027499 | 18767 | GO:0010389 | regulation of G2/M transition of mitotic cell cycle |
| rs13476961 | MGI:92531 | D3Mit60 | DNA segment, Chr 3, Massachusetts Institute of Technology 60 | DNA segment | | 85041 |  |  |
| rs6322205 | No associated gene | |  |  |  |  |  |  |
| rs13479755 | No associated gene | |  |  |  |  |  |  |
| rs6244635 | No associated gene | |  |  |  |  |  |  |
| rs13479654 | MGI:2443510 | Tacc1 | transforming, acidic coiled-coil containing protein 1 | protein coding gene | ENSMUSG00000065954 | 320165 | GO:0007049 | cell cycle |
| rs13479654 | MGI:2443510 | Tacc1 | transforming, acidic coiled-coil containing protein 1 | protein coding gene | ENSMUSG00000065954 | 320165 | GO:0051301 | cell division |
| rs13479654 | MGI:2443510 | Tacc1 | transforming, acidic coiled-coil containing protein 1 | protein coding gene | ENSMUSG00000065954 | 320165 | GO:0008283 | cell population proliferation |
| rs13479654 | MGI:2443510 | Tacc1 | transforming, acidic coiled-coil containing protein 1 | protein coding gene | ENSMUSG00000065954 | 320165 | GO:0021987 | cerebral cortex development |
| rs13479654 | MGI:2443510 | Tacc1 | transforming, acidic coiled-coil containing protein 1 | protein coding gene | ENSMUSG00000065954 | 320165 | GO:0005856 | cytoskeleton |
| rs13479654 | MGI:2443510 | Tacc1 | transforming, acidic coiled-coil containing protein 1 | protein coding gene | ENSMUSG00000065954 | 320165 | GO:0005829 | cytosol |
| rs13479654 | MGI:2443510 | Tacc1 | transforming, acidic coiled-coil containing protein 1 | protein coding gene | ENSMUSG00000065954 | 320165 | GO:0030331 | estrogen receptor binding |
| rs13479654 | MGI:2443510 | Tacc1 | transforming, acidic coiled-coil containing protein 1 | protein coding gene | ENSMUSG00000065954 | 320165 | GO:0035259 | glucocorticoid receptor binding |
| rs13479654 | MGI:2443510 | Tacc1 | transforming, acidic coiled-coil containing protein 1 | protein coding gene | ENSMUSG00000065954 | 320165 | GO:0022027 | interkinetic nuclear migration |
| rs13479654 | MGI:2443510 | Tacc1 | transforming, acidic coiled-coil containing protein 1 | protein coding gene | ENSMUSG00000065954 | 320165 | GO:0000226 | microtubule cytoskeleton organization |
| rs13479654 | MGI:2443510 | Tacc1 | transforming, acidic coiled-coil containing protein 1 | protein coding gene | ENSMUSG00000065954 | 320165 | GO:0007052 | mitotic spindle organization |
| rs13479654 | MGI:2443510 | Tacc1 | transforming, acidic coiled-coil containing protein 1 | protein coding gene | ENSMUSG00000065954 | 320165 | GO:0022008 | neurogenesis |
| rs13479654 | MGI:2443510 | Tacc1 | transforming, acidic coiled-coil containing protein 1 | protein coding gene | ENSMUSG00000065954 | 320165 | GO:0016922 | nuclear receptor binding |
| rs13479654 | MGI:2443510 | Tacc1 | transforming, acidic coiled-coil containing protein 1 | protein coding gene | ENSMUSG00000065954 | 320165 | GO:0005634 | nucleus |
| rs13479654 | MGI:2443510 | Tacc1 | transforming, acidic coiled-coil containing protein 1 | protein coding gene | ENSMUSG00000065954 | 320165 | GO:0042975 | peroxisome proliferator activated receptor binding |
| rs13479654 | MGI:2443510 | Tacc1 | transforming, acidic coiled-coil containing protein 1 | protein coding gene | ENSMUSG00000065954 | 320165 | GO:0042975 | peroxisome proliferator activated receptor binding |
| rs13479654 | MGI:2443510 | Tacc1 | transforming, acidic coiled-coil containing protein 1 | protein coding gene | ENSMUSG00000065954 | 320165 | GO:2000327 | positive regulation of nuclear receptor transcription coactivator activity |
| rs13479654 | MGI:2443510 | Tacc1 | transforming, acidic coiled-coil containing protein 1 | protein coding gene | ENSMUSG00000065954 | 320165 | GO:0019904 | protein domain specific binding |
| rs13479654 | MGI:2443510 | Tacc1 | transforming, acidic coiled-coil containing protein 1 | protein coding gene | ENSMUSG00000065954 | 320165 | GO:0032886 | regulation of microtubule-based process |
| rs13479654 | MGI:2443510 | Tacc1 | transforming, acidic coiled-coil containing protein 1 | protein coding gene | ENSMUSG00000065954 | 320165 | GO:0042974 | retinoic acid receptor binding |
| rs13479654 | MGI:2443510 | Tacc1 | transforming, acidic coiled-coil containing protein 1 | protein coding gene | ENSMUSG00000065954 | 320165 | GO:0042974 | retinoic acid receptor binding |
| rs13479654 | MGI:2443510 | Tacc1 | transforming, acidic coiled-coil containing protein 1 | protein coding gene | ENSMUSG00000065954 | 320165 | GO:0046965 | retinoid X receptor binding |
| rs13479654 | MGI:2443510 | Tacc1 | transforming, acidic coiled-coil containing protein 1 | protein coding gene | ENSMUSG00000065954 | 320165 | GO:0046965 | retinoid X receptor binding |
| rs13479654 | MGI:2443510 | Tacc1 | transforming, acidic coiled-coil containing protein 1 | protein coding gene | ENSMUSG00000065954 | 320165 | GO:0046966 | thyroid hormone receptor binding |
| rs13479654 | MGI:2443510 | Tacc1 | transforming, acidic coiled-coil containing protein 1 | protein coding gene | ENSMUSG00000065954 | 320165 | GO:0046966 | thyroid hormone receptor binding |
| rs3725706 | MGI:1289213 | D3Ertd751e | DNA segment, Chr 3, ERATO Doi 751, expressed | protein coding gene | ENSMUSG00000025766 | 73852 | GO:0008150 | biological_process |
| rs3725706 | MGI:1289213 | D3Ertd751e | DNA segment, Chr 3, ERATO Doi 751, expressed | protein coding gene | ENSMUSG00000025766 | 73852 | GO:0005575 | cellular_component |
| rs3725706 | MGI:1289213 | D3Ertd751e | DNA segment, Chr 3, ERATO Doi 751, expressed | protein coding gene | ENSMUSG00000025766 | 73852 | GO:0003674 | molecular_function |
| rs13476396 | No associated gene | |  |  |  |  |  |  |
| rs6324747 | No associated gene | |  |  |  |  |  |  |
| rs3718146 | No associated gene | |  |  |  |  |  |  |
| rs13477755 | No associated gene | |  |  |  |  |  |  |
| rs3672384 | MGI:5622948 | Gm40063 | predicted gene, 40063 | lncRNA gene | | 105244458 |  |  |
| rs13482019 | MGI:5623952 | Gm41067 | predicted gene, 41067 | lncRNA gene | | 105245639 |  |  |
| rs3694031 | MGI:5683933 | Cpgi18858 | CpG island 18858 | CpG island |  |  |  |  |
| rs3694031 | MGI:97542 | Pepd | peptidase D | protein coding gene | ENSMUSG00000063931 | 18624 | GO:0004177 | aminopeptidase activity |
| rs3694031 | MGI:97542 | Pepd | peptidase D | protein coding gene | ENSMUSG00000063931 | 18624 | GO:0030574 | collagen catabolic process |
| rs3694031 | MGI:97542 | Pepd | peptidase D | protein coding gene | ENSMUSG00000063931 | 18624 | GO:0016805 | dipeptidase activity |
| rs3694031 | MGI:97542 | Pepd | peptidase D | protein coding gene | ENSMUSG00000063931 | 18624 | GO:0016787 | hydrolase activity |
| rs3694031 | MGI:97542 | Pepd | peptidase D | protein coding gene | ENSMUSG00000063931 | 18624 | GO:0030145 | manganese ion binding |
| rs3694031 | MGI:97542 | Pepd | peptidase D | protein coding gene | ENSMUSG00000063931 | 18624 | GO:0046872 | metal ion binding |
| rs3694031 | MGI:97542 | Pepd | peptidase D | protein coding gene | ENSMUSG00000063931 | 18624 | GO:0008237 | metallopeptidase activity |
| rs3694031 | MGI:97542 | Pepd | peptidase D | protein coding gene | ENSMUSG00000063931 | 18624 | GO:0008233 | peptidase activity |
| rs3694031 | MGI:97542 | Pepd | peptidase D | protein coding gene | ENSMUSG00000063931 | 18624 | GO:0102009 | proline dipeptidase activity |
| rs3694031 | MGI:97542 | Pepd | peptidase D | protein coding gene | ENSMUSG00000063931 | 18624 | GO:0006508 | proteolysis |
| rs3663861 | No associated gene | |  |  |  |  |  |  |
| rs13482018 | MGI:102670 | Ddx4 | DEAD box helicase 4 | protein coding gene | ENSMUSG00000021758 | 13206 | GO:0016887 | ATPase activity |
| rs13482018 | MGI:102670 | Ddx4 | DEAD box helicase 4 | protein coding gene | ENSMUSG00000021758 | 13206 | GO:0005524 | ATP binding |
| rs13482018 | MGI:102670 | Ddx4 | DEAD box helicase 4 | protein coding gene | ENSMUSG00000021758 | 13206 | GO:0033391 | chromatoid body |
| rs13482018 | MGI:102670 | Ddx4 | DEAD box helicase 4 | protein coding gene | ENSMUSG00000021758 | 13206 | GO:0033391 | chromatoid body |
| rs13482018 | MGI:102670 | Ddx4 | DEAD box helicase 4 | protein coding gene | ENSMUSG00000021758 | 13206 | GO:0005737 | cytoplasm |
| rs13482018 | MGI:102670 | Ddx4 | DEAD box helicase 4 | protein coding gene | ENSMUSG00000021758 | 13206 | GO:0005737 | cytoplasm |
| rs13482018 | MGI:102670 | Ddx4 | DEAD box helicase 4 | protein coding gene | ENSMUSG00000021758 | 13206 | GO:0005737 | cytoplasm |
| rs13482018 | MGI:102670 | Ddx4 | DEAD box helicase 4 | protein coding gene | ENSMUSG00000021758 | 13206 | GO:0043046 | DNA methylation involved in gamete generation |
| rs13482018 | MGI:102670 | Ddx4 | DEAD box helicase 4 | protein coding gene | ENSMUSG00000021758 | 13206 | GO:0030317 | flagellated sperm motility |
| rs13482018 | MGI:102670 | Ddx4 | DEAD box helicase 4 | protein coding gene | ENSMUSG00000021758 | 13206 | GO:0031047 | gene silencing by RNA |
| rs13482018 | MGI:102670 | Ddx4 | DEAD box helicase 4 | protein coding gene | ENSMUSG00000021758 | 13206 | GO:0004386 | helicase activity |
| rs13482018 | MGI:102670 | Ddx4 | DEAD box helicase 4 | protein coding gene | ENSMUSG00000021758 | 13206 | GO:0016787 | hydrolase activity |
| rs13482018 | MGI:102670 | Ddx4 | DEAD box helicase 4 | protein coding gene | ENSMUSG00000021758 | 13206 | GO:0007141 | male meiosis I |
| rs13482018 | MGI:102670 | Ddx4 | DEAD box helicase 4 | protein coding gene | ENSMUSG00000021758 | 13206 | GO:0007140 | male meiotic nuclear division |
| rs13482018 | MGI:102670 | Ddx4 | DEAD box helicase 4 | protein coding gene | ENSMUSG00000021758 | 13206 | GO:0051321 | meiotic cell cycle |
| rs13482018 | MGI:102670 | Ddx4 | DEAD box helicase 4 | protein coding gene | ENSMUSG00000021758 | 13206 | GO:0007275 | multicellular organism development |
| rs13482018 | MGI:102670 | Ddx4 | DEAD box helicase 4 | protein coding gene | ENSMUSG00000021758 | 13206 | GO:0010529 | negative regulation of transposition |
| rs13482018 | MGI:102670 | Ddx4 | DEAD box helicase 4 | protein coding gene | ENSMUSG00000021758 | 13206 | GO:0003676 | nucleic acid binding |
| rs13482018 | MGI:102670 | Ddx4 | DEAD box helicase 4 | protein coding gene | ENSMUSG00000021758 | 13206 | GO:0000166 | nucleotide binding |
| rs13482018 | MGI:102670 | Ddx4 | DEAD box helicase 4 | protein coding gene | ENSMUSG00000021758 | 13206 | GO:0005634 | nucleus |
| rs13482018 | MGI:102670 | Ddx4 | DEAD box helicase 4 | protein coding gene | ENSMUSG00000021758 | 13206 | GO:0048471 | perinuclear region of cytoplasm |
| rs13482018 | MGI:102670 | Ddx4 | DEAD box helicase 4 | protein coding gene | ENSMUSG00000021758 | 13206 | GO:0048471 | perinuclear region of cytoplasm |
| rs13482018 | MGI:102670 | Ddx4 | DEAD box helicase 4 | protein coding gene | ENSMUSG00000021758 | 13206 | GO:0048471 | perinuclear region of cytoplasm |
| rs13482018 | MGI:102670 | Ddx4 | DEAD box helicase 4 | protein coding gene | ENSMUSG00000021758 | 13206 | GO:0071546 | pi-body |
| rs13482018 | MGI:102670 | Ddx4 | DEAD box helicase 4 | protein coding gene | ENSMUSG00000021758 | 13206 | GO:0071547 | piP-body |
| rs13482018 | MGI:102670 | Ddx4 | DEAD box helicase 4 | protein coding gene | ENSMUSG00000021758 | 13206 | GO:1990511 | piRNA biosynthetic process |
| rs13482018 | MGI:102670 | Ddx4 | DEAD box helicase 4 | protein coding gene | ENSMUSG00000021758 | 13206 | GO:0034587 | piRNA metabolic process |
| rs13482018 | MGI:102670 | Ddx4 | DEAD box helicase 4 | protein coding gene | ENSMUSG00000021758 | 13206 | GO:0005515 | protein binding |
| rs13482018 | MGI:102670 | Ddx4 | DEAD box helicase 4 | protein coding gene | ENSMUSG00000021758 | 13206 | GO:0032880 | regulation of protein localization |
| rs13482018 | MGI:102670 | Ddx4 | DEAD box helicase 4 | protein coding gene | ENSMUSG00000021758 | 13206 | GO:1990904 | ribonucleoprotein complex |
| rs13482018 | MGI:102670 | Ddx4 | DEAD box helicase 4 | protein coding gene | ENSMUSG00000021758 | 13206 | GO:0007283 | spermatogenesis |
| rs3719853 | MGI:5623954 | Gm41069 | predicted gene, 41069 | lncRNA gene | | 105245641 |  |  |
| rs3719311 | MGI:5683920 | Cpgi18845 | CpG island 18845 | CpG island |  |  |  |  |
| rs3719311 | MGI:3702543 | Gm12758 | predicted gene 12758 | lncRNA gene | ENSMUSG00000085105 | 100126229 | GO:0008150 | biological_process |
| rs3719311 | MGI:3702543 | Gm12758 | predicted gene 12758 | lncRNA gene | ENSMUSG00000085105 | 100126229 | GO:0005575 | cellular_component |
| rs3719311 | MGI:3702543 | Gm12758 | predicted gene 12758 | lncRNA gene | ENSMUSG00000085105 | 100126229 | GO:0003674 | molecular_function |
| rs3719311 | MGI:3649976 | Gm12762 | predicted gene 12762 | lncRNA gene | ENSMUSG00000086526 | 105247247 |  |  |
| rs3719311 | MGI:5999763 | Tssr71996 | transcription start site region 71996 | TSS region |  |  |  |  |
| rs3719311 | MGI:5999764 | Tssr71997 | transcription start site region 71997 | TSS region |  |  |  |  |
| rs3719311 | MGI:5999765 | Tssr71998 | transcription start site region 71998 | TSS region |  |  |  |  |
| rs6239372 | MGI:1916197 | Chst8 | carbohydrate sulfotransferase 8 | protein coding gene | ENSMUSG00000060402 | 68947 | GO:0016051 | carbohydrate biosynthetic process |
| rs6239372 | MGI:1916197 | Chst8 | carbohydrate sulfotransferase 8 | protein coding gene | ENSMUSG00000060402 | 68947 | GO:0005975 | carbohydrate metabolic process |
| rs6239372 | MGI:1916197 | Chst8 | carbohydrate sulfotransferase 8 | protein coding gene | ENSMUSG00000060402 | 68947 | GO:0005794 | Golgi apparatus |
| rs6239372 | MGI:1916197 | Chst8 | carbohydrate sulfotransferase 8 | protein coding gene | ENSMUSG00000060402 | 68947 | GO:0016021 | integral component of membrane |
| rs6239372 | MGI:1916197 | Chst8 | carbohydrate sulfotransferase 8 | protein coding gene | ENSMUSG00000060402 | 68947 | GO:0016020 | membrane |
| rs6239372 | MGI:1916197 | Chst8 | carbohydrate sulfotransferase 8 | protein coding gene | ENSMUSG00000060402 | 68947 | GO:0001537 | N-acetylgalactosamine 4-O-sulfotransferase activity |
| rs6239372 | MGI:1916197 | Chst8 | carbohydrate sulfotransferase 8 | protein coding gene | ENSMUSG00000060402 | 68947 | GO:0001537 | N-acetylgalactosamine 4-O-sulfotransferase activity |
| rs6239372 | MGI:1916197 | Chst8 | carbohydrate sulfotransferase 8 | protein coding gene | ENSMUSG00000060402 | 68947 | GO:0016486 | peptide hormone processing |
| rs6239372 | MGI:1916197 | Chst8 | carbohydrate sulfotransferase 8 | protein coding gene | ENSMUSG00000060402 | 68947 | GO:0030166 | proteoglycan biosynthetic process |
| rs6239372 | MGI:1916197 | Chst8 | carbohydrate sulfotransferase 8 | protein coding gene | ENSMUSG00000060402 | 68947 | GO:0030166 | proteoglycan biosynthetic process |
| rs6239372 | MGI:1916197 | Chst8 | carbohydrate sulfotransferase 8 | protein coding gene | ENSMUSG00000060402 | 68947 | GO:0006790 | sulfur compound metabolic process |
| rs6239372 | MGI:1916197 | Chst8 | carbohydrate sulfotransferase 8 | protein coding gene | ENSMUSG00000060402 | 68947 | GO:0016740 | transferase activity |
| rs3694208 | MGI:2443181 | Dlgap2 | DLG associated protein 2 | protein coding gene | ENSMUSG00000047495 | 244310 | GO:0030054 | cell junction |
| rs3694208 | MGI:2443181 | Dlgap2 | DLG associated protein 2 | protein coding gene | ENSMUSG00000047495 | 244310 | GO:0030425 | dendrite |
| rs3694208 | MGI:2443181 | Dlgap2 | DLG associated protein 2 | protein coding gene | ENSMUSG00000047495 | 244310 | GO:0043197 | dendritic spine |
| rs3694208 | MGI:2443181 | Dlgap2 | DLG associated protein 2 | protein coding gene | ENSMUSG00000047495 | 244310 | GO:0016020 | membrane |
| rs3694208 | MGI:2443181 | Dlgap2 | DLG associated protein 2 | protein coding gene | ENSMUSG00000047495 | 244310 | GO:0007270 | neuron-neuron synaptic transmission |
| rs3694208 | MGI:2443181 | Dlgap2 | DLG associated protein 2 | protein coding gene | ENSMUSG00000047495 | 244310 | GO:0005886 | plasma membrane |
| rs3694208 | MGI:2443181 | Dlgap2 | DLG associated protein 2 | protein coding gene | ENSMUSG00000047495 | 244310 | GO:0019904 | protein domain specific binding |
| rs3694208 | MGI:2443181 | Dlgap2 | DLG associated protein 2 | protein coding gene | ENSMUSG00000047495 | 244310 | GO:0098962 | regulation of postsynaptic neurotransmitter receptor activity |
| rs3694208 | MGI:2443181 | Dlgap2 | DLG associated protein 2 | protein coding gene | ENSMUSG00000047495 | 244310 | GO:0098962 | regulation of postsynaptic neurotransmitter receptor activity |
| rs3694208 | MGI:2443181 | Dlgap2 | DLG associated protein 2 | protein coding gene | ENSMUSG00000047495 | 244310 | GO:0023052 | signaling |
| rs3694208 | MGI:2443181 | Dlgap2 | DLG associated protein 2 | protein coding gene | ENSMUSG00000047495 | 244310 | GO:0045202 | synapse |
| rs3679439 | No associated gene | |  |  |  |  |  |  |
| rs13482016 | No associated gene | |  |  |  |  |  |  |
| rs3708535 | MGI:1927222 | Diaph3 | diaphanous related formin 3 | protein coding gene | ENSMUSG00000022021 | 56419 | GO:0003779 | actin binding |
| rs3708535 | MGI:1927222 | Diaph3 | diaphanous related formin 3 | protein coding gene | ENSMUSG00000022021 | 56419 | GO:0030036 | actin cytoskeleton organization |
| rs3708535 | MGI:1927222 | Diaph3 | diaphanous related formin 3 | protein coding gene | ENSMUSG00000022021 | 56419 | GO:0030041 | actin filament polymerization |
| rs3708535 | MGI:1927222 | Diaph3 | diaphanous related formin 3 | protein coding gene | ENSMUSG00000022021 | 56419 | GO:0016043 | cellular component organization |
| rs3708535 | MGI:1927222 | Diaph3 | diaphanous related formin 3 | protein coding gene | ENSMUSG00000022021 | 56419 | GO:0005737 | cytoplasm |
| rs3708535 | MGI:1927222 | Diaph3 | diaphanous related formin 3 | protein coding gene | ENSMUSG00000022021 | 56419 | GO:0005737 | cytoplasm |
| rs3708535 | MGI:1927222 | Diaph3 | diaphanous related formin 3 | protein coding gene | ENSMUSG00000022021 | 56419 | GO:0007010 | cytoskeleton organization |
| rs3708535 | MGI:1927222 | Diaph3 | diaphanous related formin 3 | protein coding gene | ENSMUSG00000022021 | 56419 | GO:0005634 | nucleus |
| rs3708535 | MGI:1927222 | Diaph3 | diaphanous related formin 3 | protein coding gene | ENSMUSG00000022021 | 56419 | GO:0005634 | nucleus |
| rs3708535 | MGI:1927222 | Diaph3 | diaphanous related formin 3 | protein coding gene | ENSMUSG00000022021 | 56419 | GO:0005515 | protein binding |
| rs3708535 | MGI:1927222 | Diaph3 | diaphanous related formin 3 | protein coding gene | ENSMUSG00000022021 | 56419 | GO:0017048 | Rho GTPase binding |
| rs13479188 | MGI:1289234 | Rhpn2 | rhophilin, Rho GTPase binding protein 2 | protein coding gene | ENSMUSG00000030494 | 52428 | GO:0005737 | cytoplasm |
| rs13479188 | MGI:1289234 | Rhpn2 | rhophilin, Rho GTPase binding protein 2 | protein coding gene | ENSMUSG00000030494 | 52428 | GO:0003674 | molecular_function |
| rs13479188 | MGI:1289234 | Rhpn2 | rhophilin, Rho GTPase binding protein 2 | protein coding gene | ENSMUSG00000030494 | 52428 | GO:0007165 | signal transduction |
| rs3725510 | No associated gene | |  |  |  |  |  |  |
| rs13480302 | No associated gene | |  |  |  |  |  |  |
| rs13480638 | MGI:3036232 | 4833431D13Rik | RIKEN cDNA 4833431D13 gene | lncRNA gene | | 268307 |  |  |
| rs13480638 | MGI:88026 | Ank3 | ankyrin 3, epithelial | protein coding gene | ENSMUSG00000069601 | 11735 | GO:0030424 | axon |
| rs13480638 | MGI:88026 | Ank3 | ankyrin 3, epithelial | protein coding gene | ENSMUSG00000069601 | 11735 | GO:0030424 | axon |
| rs13480638 | MGI:88026 | Ank3 | ankyrin 3, epithelial | protein coding gene | ENSMUSG00000069601 | 11735 | GO:0007411 | axon guidance |
| rs13480638 | MGI:88026 | Ank3 | ankyrin 3, epithelial | protein coding gene | ENSMUSG00000069601 | 11735 | GO:0043194 | axon initial segment |
| rs13480638 | MGI:88026 | Ank3 | ankyrin 3, epithelial | protein coding gene | ENSMUSG00000069601 | 11735 | GO:0043194 | axon initial segment |
| rs13480638 | MGI:88026 | Ank3 | ankyrin 3, epithelial | protein coding gene | ENSMUSG00000069601 | 11735 | GO:0043194 | axon initial segment |
| rs13480638 | MGI:88026 | Ank3 | ankyrin 3, epithelial | protein coding gene | ENSMUSG00000069601 | 11735 | GO:0007409 | axonogenesis |
| rs13480638 | MGI:88026 | Ank3 | ankyrin 3, epithelial | protein coding gene | ENSMUSG00000069601 | 11735 | GO:0009925 | basal plasma membrane |
| rs13480638 | MGI:88026 | Ank3 | ankyrin 3, epithelial | protein coding gene | ENSMUSG00000069601 | 11735 | GO:0016323 | basolateral plasma membrane |
| rs13480638 | MGI:88026 | Ank3 | ankyrin 3, epithelial | protein coding gene | ENSMUSG00000069601 | 11735 | GO:0005923 | bicellular tight junction |
| rs13480638 | MGI:88026 | Ank3 | ankyrin 3, epithelial | protein coding gene | ENSMUSG00000069601 | 11735 | GO:0045296 | cadherin binding |
| rs13480638 | MGI:88026 | Ank3 | ankyrin 3, epithelial | protein coding gene | ENSMUSG00000069601 | 11735 | GO:0030054 | cell junction |
| rs13480638 | MGI:88026 | Ank3 | ankyrin 3, epithelial | protein coding gene | ENSMUSG00000069601 | 11735 | GO:0042995 | cell projection |
| rs13480638 | MGI:88026 | Ank3 | ankyrin 3, epithelial | protein coding gene | ENSMUSG00000069601 | 11735 | GO:0009986 | cell surface |
| rs13480638 | MGI:88026 | Ank3 | ankyrin 3, epithelial | protein coding gene | ENSMUSG00000069601 | 11735 | GO:0071286 | cellular response to magnesium ion |
| rs13480638 | MGI:88026 | Ank3 | ankyrin 3, epithelial | protein coding gene | ENSMUSG00000069601 | 11735 | GO:0045162 | clustering of voltage-gated sodium channels |
| rs13480638 | MGI:88026 | Ank3 | ankyrin 3, epithelial | protein coding gene | ENSMUSG00000069601 | 11735 | GO:0005737 | cytoplasm |
| rs13480638 | MGI:88026 | Ank3 | ankyrin 3, epithelial | protein coding gene | ENSMUSG00000069601 | 11735 | GO:0008092 | cytoskeletal protein binding |
| rs13480638 | MGI:88026 | Ank3 | ankyrin 3, epithelial | protein coding gene | ENSMUSG00000069601 | 11735 | GO:0005856 | cytoskeleton |
| rs13480638 | MGI:88026 | Ank3 | ankyrin 3, epithelial | protein coding gene | ENSMUSG00000069601 | 11735 | GO:0030425 | dendrite |
| rs13480638 | MGI:88026 | Ank3 | ankyrin 3, epithelial | protein coding gene | ENSMUSG00000069601 | 11735 | GO:0045184 | establishment of protein localization |
| rs13480638 | MGI:88026 | Ank3 | ankyrin 3, epithelial | protein coding gene | ENSMUSG00000069601 | 11735 | GO:0043001 | Golgi to plasma membrane protein transport |
| rs13480638 | MGI:88026 | Ank3 | ankyrin 3, epithelial | protein coding gene | ENSMUSG00000069601 | 11735 | GO:0014704 | intercalated disc |
| rs13480638 | MGI:88026 | Ank3 | ankyrin 3, epithelial | protein coding gene | ENSMUSG00000069601 | 11735 | GO:0014704 | intercalated disc |
| rs13480638 | MGI:88026 | Ank3 | ankyrin 3, epithelial | protein coding gene | ENSMUSG00000069601 | 11735 | GO:0044325 | ion channel binding |
| rs13480638 | MGI:88026 | Ank3 | ankyrin 3, epithelial | protein coding gene | ENSMUSG00000069601 | 11735 | GO:0044325 | ion channel binding |
| rs13480638 | MGI:88026 | Ank3 | ankyrin 3, epithelial | protein coding gene | ENSMUSG00000069601 | 11735 | GO:0016328 | lateral plasma membrane |
| rs13480638 | MGI:88026 | Ank3 | ankyrin 3, epithelial | protein coding gene | ENSMUSG00000069601 | 11735 | GO:0005764 | lysosome |
| rs13480638 | MGI:88026 | Ank3 | ankyrin 3, epithelial | protein coding gene | ENSMUSG00000069601 | 11735 | GO:0010960 | magnesium ion homeostasis |
| rs13480638 | MGI:88026 | Ank3 | ankyrin 3, epithelial | protein coding gene | ENSMUSG00000069601 | 11735 | GO:0072660 | maintenance of protein location in plasma membrane |
| rs13480638 | MGI:88026 | Ank3 | ankyrin 3, epithelial | protein coding gene | ENSMUSG00000069601 | 11735 | GO:0016020 | membrane |
| rs13480638 | MGI:88026 | Ank3 | ankyrin 3, epithelial | protein coding gene | ENSMUSG00000069601 | 11735 | GO:0071709 | membrane assembly |
| rs13480638 | MGI:88026 | Ank3 | ankyrin 3, epithelial | protein coding gene | ENSMUSG00000069601 | 11735 | GO:0000281 | mitotic cytokinesis |
| rs13480638 | MGI:88026 | Ank3 | ankyrin 3, epithelial | protein coding gene | ENSMUSG00000069601 | 11735 | GO:1902260 | negative regulation of delayed rectifier potassium channel activity |
| rs13480638 | MGI:88026 | Ank3 | ankyrin 3, epithelial | protein coding gene | ENSMUSG00000069601 | 11735 | GO:0031594 | neuromuscular junction |
| rs13480638 | MGI:88026 | Ank3 | ankyrin 3, epithelial | protein coding gene | ENSMUSG00000069601 | 11735 | GO:0019228 | neuronal action potential |
| rs13480638 | MGI:88026 | Ank3 | ankyrin 3, epithelial | protein coding gene | ENSMUSG00000069601 | 11735 | GO:0043005 | neuron projection |
| rs13480638 | MGI:88026 | Ank3 | ankyrin 3, epithelial | protein coding gene | ENSMUSG00000069601 | 11735 | GO:0033268 | node of Ranvier |
| rs13480638 | MGI:88026 | Ank3 | ankyrin 3, epithelial | protein coding gene | ENSMUSG00000069601 | 11735 | GO:0033268 | node of Ranvier |
| rs13480638 | MGI:88026 | Ank3 | ankyrin 3, epithelial | protein coding gene | ENSMUSG00000069601 | 11735 | GO:0033270 | paranode region of axon |
| rs13480638 | MGI:88026 | Ank3 | ankyrin 3, epithelial | protein coding gene | ENSMUSG00000069601 | 11735 | GO:0140031 | phosphorylation-dependent protein binding |
| rs13480638 | MGI:88026 | Ank3 | ankyrin 3, epithelial | protein coding gene | ENSMUSG00000069601 | 11735 | GO:0005886 | plasma membrane |
| rs13480638 | MGI:88026 | Ank3 | ankyrin 3, epithelial | protein coding gene | ENSMUSG00000069601 | 11735 | GO:0007009 | plasma membrane organization |
| rs13480638 | MGI:88026 | Ank3 | ankyrin 3, epithelial | protein coding gene | ENSMUSG00000069601 | 11735 | GO:0045760 | positive regulation of action potential |
| rs13480638 | MGI:88026 | Ank3 | ankyrin 3, epithelial | protein coding gene | ENSMUSG00000069601 | 11735 | GO:2001259 | positive regulation of cation channel activity |
| rs13480638 | MGI:88026 | Ank3 | ankyrin 3, epithelial | protein coding gene | ENSMUSG00000069601 | 11735 | GO:0010650 | positive regulation of cell communication by electrical coupling |
| rs13480638 | MGI:88026 | Ank3 | ankyrin 3, epithelial | protein coding gene | ENSMUSG00000069601 | 11735 | GO:0010628 | positive regulation of gene expression |
| rs13480638 | MGI:88026 | Ank3 | ankyrin 3, epithelial | protein coding gene | ENSMUSG00000069601 | 11735 | GO:0034112 | positive regulation of homotypic cell-cell adhesion |
| rs13480638 | MGI:88026 | Ank3 | ankyrin 3, epithelial | protein coding gene | ENSMUSG00000069601 | 11735 | GO:1900827 | positive regulation of membrane depolarization during cardiac muscle cell action potential |
| rs13480638 | MGI:88026 | Ank3 | ankyrin 3, epithelial | protein coding gene | ENSMUSG00000069601 | 11735 | GO:0045838 | positive regulation of membrane potential |
| rs13480638 | MGI:88026 | Ank3 | ankyrin 3, epithelial | protein coding gene | ENSMUSG00000069601 | 11735 | GO:0090314 | positive regulation of protein targeting to membrane |
| rs13480638 | MGI:88026 | Ank3 | ankyrin 3, epithelial | protein coding gene | ENSMUSG00000069601 | 11735 | GO:2000651 | positive regulation of sodium ion transmembrane transporter activity |
| rs13480638 | MGI:88026 | Ank3 | ankyrin 3, epithelial | protein coding gene | ENSMUSG00000069601 | 11735 | GO:0010765 | positive regulation of sodium ion transport |
| rs13480638 | MGI:88026 | Ank3 | ankyrin 3, epithelial | protein coding gene | ENSMUSG00000069601 | 11735 | GO:0045211 | postsynaptic membrane |
| rs13480638 | MGI:88026 | Ank3 | ankyrin 3, epithelial | protein coding gene | ENSMUSG00000069601 | 11735 | GO:0005515 | protein binding |
| rs13480638 | MGI:88026 | Ank3 | ankyrin 3, epithelial | protein coding gene | ENSMUSG00000069601 | 11735 | GO:0099612 | protein localization to axon |
| rs13480638 | MGI:88026 | Ank3 | ankyrin 3, epithelial | protein coding gene | ENSMUSG00000069601 | 11735 | GO:0099612 | protein localization to axon |
| rs13480638 | MGI:88026 | Ank3 | ankyrin 3, epithelial | protein coding gene | ENSMUSG00000069601 | 11735 | GO:0072659 | protein localization to plasma membrane |
| rs13480638 | MGI:88026 | Ank3 | ankyrin 3, epithelial | protein coding gene | ENSMUSG00000069601 | 11735 | GO:0072659 | protein localization to plasma membrane |
| rs13480638 | MGI:88026 | Ank3 | ankyrin 3, epithelial | protein coding gene | ENSMUSG00000069601 | 11735 | GO:0072659 | protein localization to plasma membrane |
| rs13480638 | MGI:88026 | Ank3 | ankyrin 3, epithelial | protein coding gene | ENSMUSG00000069601 | 11735 | GO:0030674 | protein-macromolecule adaptor activity |
| rs13480638 | MGI:88026 | Ank3 | ankyrin 3, epithelial | protein coding gene | ENSMUSG00000069601 | 11735 | GO:0043266 | regulation of potassium ion transport |
| rs13480638 | MGI:88026 | Ank3 | ankyrin 3, epithelial | protein coding gene | ENSMUSG00000069601 | 11735 | GO:0042383 | sarcolemma |
| rs13480638 | MGI:88026 | Ank3 | ankyrin 3, epithelial | protein coding gene | ENSMUSG00000069601 | 11735 | GO:0016529 | sarcoplasmic reticulum |
| rs13480638 | MGI:88026 | Ank3 | ankyrin 3, epithelial | protein coding gene | ENSMUSG00000069601 | 11735 | GO:0007165 | signal transduction |
| rs13480638 | MGI:88026 | Ank3 | ankyrin 3, epithelial | protein coding gene | ENSMUSG00000069601 | 11735 | GO:0014731 | spectrin-associated cytoskeleton |
| rs13480638 | MGI:88026 | Ank3 | ankyrin 3, epithelial | protein coding gene | ENSMUSG00000069601 | 11735 | GO:0030507 | spectrin binding |
| rs13480638 | MGI:88026 | Ank3 | ankyrin 3, epithelial | protein coding gene | ENSMUSG00000069601 | 11735 | GO:0005200 | structural constituent of cytoskeleton |
| rs13480638 | MGI:88026 | Ank3 | ankyrin 3, epithelial | protein coding gene | ENSMUSG00000069601 | 11735 | GO:0045202 | synapse |
| rs13480638 | MGI:88026 | Ank3 | ankyrin 3, epithelial | protein coding gene | ENSMUSG00000069601 | 11735 | GO:0050808 | synapse organization |
| rs13480638 | MGI:88026 | Ank3 | ankyrin 3, epithelial | protein coding gene | ENSMUSG00000069601 | 11735 | GO:0030315 | T-tubule |
| rs13480638 | MGI:88026 | Ank3 | ankyrin 3, epithelial | protein coding gene | ENSMUSG00000069601 | 11735 | GO:0030018 | Z disc |
| rs4230072 | MGI:1918839 | Slc38a9 | solute carrier family 38, member 9 | protein coding gene | ENSMUSG00000047789 | 268706 | GO:0003333 | amino acid transmembrane transport |
| rs4230072 | MGI:1918839 | Slc38a9 | solute carrier family 38, member 9 | protein coding gene | ENSMUSG00000047789 | 268706 | GO:0003333 | amino acid transmembrane transport |
| rs4230072 | MGI:1918839 | Slc38a9 | solute carrier family 38, member 9 | protein coding gene | ENSMUSG00000047789 | 268706 | GO:0015171 | amino acid transmembrane transporter activity |
| rs4230072 | MGI:1918839 | Slc38a9 | solute carrier family 38, member 9 | protein coding gene | ENSMUSG00000047789 | 268706 | GO:0006865 | amino acid transport |
| rs4230072 | MGI:1918839 | Slc38a9 | solute carrier family 38, member 9 | protein coding gene | ENSMUSG00000047789 | 268706 | GO:0071230 | cellular response to amino acid stimulus |
| rs4230072 | MGI:1918839 | Slc38a9 | solute carrier family 38, member 9 | protein coding gene | ENSMUSG00000047789 | 268706 | GO:0005768 | endosome |
| rs4230072 | MGI:1918839 | Slc38a9 | solute carrier family 38, member 9 | protein coding gene | ENSMUSG00000047789 | 268706 | GO:1905103 | integral component of lysosomal membrane |
| rs4230072 | MGI:1918839 | Slc38a9 | solute carrier family 38, member 9 | protein coding gene | ENSMUSG00000047789 | 268706 | GO:0016021 | integral component of membrane |
| rs4230072 | MGI:1918839 | Slc38a9 | solute carrier family 38, member 9 | protein coding gene | ENSMUSG00000047789 | 268706 | GO:0061459 | L-arginine transmembrane transporter activity |
| rs4230072 | MGI:1918839 | Slc38a9 | solute carrier family 38, member 9 | protein coding gene | ENSMUSG00000047789 | 268706 | GO:0005770 | late endosome |
| rs4230072 | MGI:1918839 | Slc38a9 | solute carrier family 38, member 9 | protein coding gene | ENSMUSG00000047789 | 268706 | GO:0015190 | L-leucine transmembrane transporter activity |
| rs4230072 | MGI:1918839 | Slc38a9 | solute carrier family 38, member 9 | protein coding gene | ENSMUSG00000047789 | 268706 | GO:0005765 | lysosomal membrane |
| rs4230072 | MGI:1918839 | Slc38a9 | solute carrier family 38, member 9 | protein coding gene | ENSMUSG00000047789 | 268706 | GO:0005764 | lysosome |
| rs4230072 | MGI:1918839 | Slc38a9 | solute carrier family 38, member 9 | protein coding gene | ENSMUSG00000047789 | 268706 | GO:0016020 | membrane |
| rs4230072 | MGI:1918839 | Slc38a9 | solute carrier family 38, member 9 | protein coding gene | ENSMUSG00000047789 | 268706 | GO:0046872 | metal ion binding |
| rs4230072 | MGI:1918839 | Slc38a9 | solute carrier family 38, member 9 | protein coding gene | ENSMUSG00000047789 | 268706 | GO:0032008 | positive regulation of TOR signaling |
| rs4230072 | MGI:1918839 | Slc38a9 | solute carrier family 38, member 9 | protein coding gene | ENSMUSG00000047789 | 268706 | GO:0071986 | Ragulator complex |
| rs13482148 | MGI:1925040 | A930006J02Rik | RIKEN cDNA A930006J02 gene | lncRNA gene | | 77790 |  |  |
| rs13482148 | MGI:109515 | Sftpd | surfactant associated protein D | protein coding gene | ENSMUSG00000021795 | 20390 | GO:0030246 | carbohydrate binding |
| rs13482148 | MGI:109515 | Sftpd | surfactant associated protein D | protein coding gene | ENSMUSG00000021795 | 20390 | GO:0062023 | collagen-containing extracellular matrix |
| rs13482148 | MGI:109515 | Sftpd | surfactant associated protein D | protein coding gene | ENSMUSG00000021795 | 20390 | GO:0005581 | collagen trimer |
| rs13482148 | MGI:109515 | Sftpd | surfactant associated protein D | protein coding gene | ENSMUSG00000021795 | 20390 | GO:0005737 | cytoplasm |
| rs13482148 | MGI:109515 | Sftpd | surfactant associated protein D | protein coding gene | ENSMUSG00000021795 | 20390 | GO:0031410 | cytoplasmic vesicle |
| rs13482148 | MGI:109515 | Sftpd | surfactant associated protein D | protein coding gene | ENSMUSG00000021795 | 20390 | GO:0005576 | extracellular region |
| rs13482148 | MGI:109515 | Sftpd | surfactant associated protein D | protein coding gene | ENSMUSG00000021795 | 20390 | GO:0005615 | extracellular space |
| rs13482148 | MGI:109515 | Sftpd | surfactant associated protein D | protein coding gene | ENSMUSG00000021795 | 20390 | GO:0042802 | identical protein binding |
| rs13482148 | MGI:109515 | Sftpd | surfactant associated protein D | protein coding gene | ENSMUSG00000021795 | 20390 | GO:0042802 | identical protein binding |
| rs13482148 | MGI:109515 | Sftpd | surfactant associated protein D | protein coding gene | ENSMUSG00000021795 | 20390 | GO:0002376 | immune system process |
| rs13482148 | MGI:109515 | Sftpd | surfactant associated protein D | protein coding gene | ENSMUSG00000021795 | 20390 | GO:0043152 | induction of bacterial agglutination |
| rs13482148 | MGI:109515 | Sftpd | surfactant associated protein D | protein coding gene | ENSMUSG00000021795 | 20390 | GO:0045087 | innate immune response |
| rs13482148 | MGI:109515 | Sftpd | surfactant associated protein D | protein coding gene | ENSMUSG00000021795 | 20390 | GO:0001530 | lipopolysaccharide binding |
| rs13482148 | MGI:109515 | Sftpd | surfactant associated protein D | protein coding gene | ENSMUSG00000021795 | 20390 | GO:0048286 | lung alveolus development |
| rs13482148 | MGI:109515 | Sftpd | surfactant associated protein D | protein coding gene | ENSMUSG00000021795 | 20390 | GO:0048029 | monosaccharide binding |
| rs13482148 | MGI:109515 | Sftpd | surfactant associated protein D | protein coding gene | ENSMUSG00000021795 | 20390 | GO:0005771 | multivesicular body |
| rs13482148 | MGI:109515 | Sftpd | surfactant associated protein D | protein coding gene | ENSMUSG00000021795 | 20390 | GO:0052405 | negative regulation by host of symbiont molecular function |
| rs13482148 | MGI:109515 | Sftpd | surfactant associated protein D | protein coding gene | ENSMUSG00000021795 | 20390 | GO:0032703 | negative regulation of interleukin-2 production |
| rs13482148 | MGI:109515 | Sftpd | surfactant associated protein D | protein coding gene | ENSMUSG00000021795 | 20390 | GO:0050765 | negative regulation of phagocytosis |
| rs13482148 | MGI:109515 | Sftpd | surfactant associated protein D | protein coding gene | ENSMUSG00000021795 | 20390 | GO:0042130 | negative regulation of T cell proliferation |
| rs13482148 | MGI:109515 | Sftpd | surfactant associated protein D | protein coding gene | ENSMUSG00000021795 | 20390 | GO:0008228 | opsonization |
| rs13482148 | MGI:109515 | Sftpd | surfactant associated protein D | protein coding gene | ENSMUSG00000021795 | 20390 | GO:0050766 | positive regulation of phagocytosis |
| rs13482148 | MGI:109515 | Sftpd | surfactant associated protein D | protein coding gene | ENSMUSG00000021795 | 20390 | GO:1905226 | regulation of adhesion of symbiont to host epithelial cell |
| rs13482148 | MGI:109515 | Sftpd | surfactant associated protein D | protein coding gene | ENSMUSG00000021795 | 20390 | GO:0050828 | regulation of liquid surface tension |
| rs13482148 | MGI:109515 | Sftpd | surfactant associated protein D | protein coding gene | ENSMUSG00000021795 | 20390 | GO:0007585 | respiratory gaseous exchange by respiratory system |
| rs13482148 | MGI:109515 | Sftpd | surfactant associated protein D | protein coding gene | ENSMUSG00000021795 | 20390 | GO:0005791 | rough endoplasmic reticulum |
| rs13482148 | MGI:109515 | Sftpd | surfactant associated protein D | protein coding gene | ENSMUSG00000021795 | 20390 | GO:0043129 | surfactant homeostasis |
| rs13482148 | MGI:6055979 | Tssr128212 | transcription start site region 128212 | TSS region |  |  |  |  |
| rs13482148 | MGI:6055980 | Tssr128213 | transcription start site region 128213 | TSS region |  |  |  |  |
| rs13482148 | MGI:6055981 | Tssr128214 | transcription start site region 128214 | TSS region |  |  |  |  |
| rs3699140 | No associated gene | |  |  |  |  |  |  |
| rs13483571 | MGI:107930 | Pip5k1b | phosphatidylinositol-4-phosphate 5-kinase, type 1 beta | protein coding gene | ENSMUSG00000024867 | 18719 | GO:0016308 | 1-phosphatidylinositol-4-phosphate 5-kinase activity |
| rs13483571 | MGI:107930 | Pip5k1b | phosphatidylinositol-4-phosphate 5-kinase, type 1 beta | protein coding gene | ENSMUSG00000024867 | 18719 | GO:0016308 | 1-phosphatidylinositol-4-phosphate 5-kinase activity |
| rs13483571 | MGI:107930 | Pip5k1b | phosphatidylinositol-4-phosphate 5-kinase, type 1 beta | protein coding gene | ENSMUSG00000024867 | 18719 | GO:0005524 | ATP binding |
| rs13483571 | MGI:107930 | Pip5k1b | phosphatidylinositol-4-phosphate 5-kinase, type 1 beta | protein coding gene | ENSMUSG00000024867 | 18719 | GO:0005829 | cytosol |
| rs13483571 | MGI:107930 | Pip5k1b | phosphatidylinositol-4-phosphate 5-kinase, type 1 beta | protein coding gene | ENSMUSG00000024867 | 18719 | GO:0016301 | kinase activity |
| rs13483571 | MGI:107930 | Pip5k1b | phosphatidylinositol-4-phosphate 5-kinase, type 1 beta | protein coding gene | ENSMUSG00000024867 | 18719 | GO:0016020 | membrane |
| rs13483571 | MGI:107930 | Pip5k1b | phosphatidylinositol-4-phosphate 5-kinase, type 1 beta | protein coding gene | ENSMUSG00000024867 | 18719 | GO:0000166 | nucleotide binding |
| rs13483571 | MGI:107930 | Pip5k1b | phosphatidylinositol-4-phosphate 5-kinase, type 1 beta | protein coding gene | ENSMUSG00000024867 | 18719 | GO:0006661 | phosphatidylinositol biosynthetic process |
| rs13483571 | MGI:107930 | Pip5k1b | phosphatidylinositol-4-phosphate 5-kinase, type 1 beta | protein coding gene | ENSMUSG00000024867 | 18719 | GO:0046488 | phosphatidylinositol metabolic process |
| rs13483571 | MGI:107930 | Pip5k1b | phosphatidylinositol-4-phosphate 5-kinase, type 1 beta | protein coding gene | ENSMUSG00000024867 | 18719 | GO:0016307 | phosphatidylinositol phosphate kinase activity |
| rs13483571 | MGI:107930 | Pip5k1b | phosphatidylinositol-4-phosphate 5-kinase, type 1 beta | protein coding gene | ENSMUSG00000024867 | 18719 | GO:0016310 | phosphorylation |
| rs13483571 | MGI:107930 | Pip5k1b | phosphatidylinositol-4-phosphate 5-kinase, type 1 beta | protein coding gene | ENSMUSG00000024867 | 18719 | GO:0005515 | protein binding |
| rs13483571 | MGI:107930 | Pip5k1b | phosphatidylinositol-4-phosphate 5-kinase, type 1 beta | protein coding gene | ENSMUSG00000024867 | 18719 | GO:0016740 | transferase activity |
| rs13483571 | MGI:107930 | Pip5k1b | phosphatidylinositol-4-phosphate 5-kinase, type 1 beta | protein coding gene | ENSMUSG00000024867 | 18719 | GO:0001931 | uropod |
| rs13483571 | MGI:6086432 | Tssr158665 | transcription start site region 158665 | TSS region |  |  |  |  |
| rs3679568 | No associated gene | |  |  |  |  |  |  |
| rs13479566 | MGI:96832 | Lsp1 | lymphocyte specific 1 | protein coding gene | ENSMUSG00000018819 | 16985 | GO:0003779 | actin binding |
| rs13479566 | MGI:96832 | Lsp1 | lymphocyte specific 1 | protein coding gene | ENSMUSG00000018819 | 16985 | GO:0006915 | apoptotic process |
| rs13479566 | MGI:96832 | Lsp1 | lymphocyte specific 1 | protein coding gene | ENSMUSG00000018819 | 16985 | GO:0098761 | cellular response to interleukin-7 |
| rs13479566 | MGI:96832 | Lsp1 | lymphocyte specific 1 | protein coding gene | ENSMUSG00000018819 | 16985 | GO:0006935 | chemotaxis |
| rs13479566 | MGI:96832 | Lsp1 | lymphocyte specific 1 | protein coding gene | ENSMUSG00000018819 | 16985 | GO:0007010 | cytoskeleton organization |
| rs13479566 | MGI:96832 | Lsp1 | lymphocyte specific 1 | protein coding gene | ENSMUSG00000018819 | 16985 | GO:0006952 | defense response |
| rs13479566 | MGI:96832 | Lsp1 | lymphocyte specific 1 | protein coding gene | ENSMUSG00000018819 | 16985 | GO:0016020 | membrane |
| rs13479566 | MGI:96832 | Lsp1 | lymphocyte specific 1 | protein coding gene | ENSMUSG00000018819 | 16985 | GO:0005886 | plasma membrane |
| rs13479566 | MGI:96832 | Lsp1 | lymphocyte specific 1 | protein coding gene | ENSMUSG00000018819 | 16985 | GO:0007165 | signal transduction |
| rs13479566 | MGI:5930221 | Tssr2454 | transcription start site region 2454 | TSS region |  |  |  |  |
| rs13479566 | MGI:5997937 | Tssr70170 | transcription start site region 70170 | TSS region |  |  |  |  |
| rs13479566 | MGI:5997938 | Tssr70171 | transcription start site region 70171 | TSS region |  |  |  |  |
| rs4140004 | MGI:97851 | Slc20a2 | solute carrier family 20, member 2 | protein coding gene | ENSMUSG00000037656 | 20516 | GO:0016021 | integral component of membrane |
| rs4140004 | MGI:97851 | Slc20a2 | solute carrier family 20, member 2 | protein coding gene | ENSMUSG00000037656 | 20516 | GO:0006811 | ion transport |
| rs4140004 | MGI:97851 | Slc20a2 | solute carrier family 20, member 2 | protein coding gene | ENSMUSG00000037656 | 20516 | GO:0016020 | membrane |
| rs4140004 | MGI:97851 | Slc20a2 | solute carrier family 20, member 2 | protein coding gene | ENSMUSG00000037656 | 20516 | GO:0006817 | phosphate ion transport |
| rs4140004 | MGI:97851 | Slc20a2 | solute carrier family 20, member 2 | protein coding gene | ENSMUSG00000037656 | 20516 | GO:0005886 | plasma membrane |
| rs4140004 | MGI:97851 | Slc20a2 | solute carrier family 20, member 2 | protein coding gene | ENSMUSG00000037656 | 20516 | GO:0006814 | sodium ion transport |
| rs4140004 | MGI:97851 | Slc20a2 | solute carrier family 20, member 2 | protein coding gene | ENSMUSG00000037656 | 20516 | GO:0015293 | symporter activity |
| rs4140004 | MGI:97851 | Slc20a2 | solute carrier family 20, member 2 | protein coding gene | ENSMUSG00000037656 | 20516 | GO:0016032 | viral process |
| rs13459102 | MGI:1916992 | Tm2d2 | TM2 domain containing 2 | protein coding gene | ENSMUSG00000031556 | 69742 | GO:0008150 | biological_process |
| rs13459102 | MGI:1916992 | Tm2d2 | TM2 domain containing 2 | protein coding gene | ENSMUSG00000031556 | 69742 | GO:0016021 | integral component of membrane |
| rs13459102 | MGI:1916992 | Tm2d2 | TM2 domain containing 2 | protein coding gene | ENSMUSG00000031556 | 69742 | GO:0016020 | membrane |
| rs13459102 | MGI:1916992 | Tm2d2 | TM2 domain containing 2 | protein coding gene | ENSMUSG00000031556 | 69742 | GO:0003674 | molecular_function |
| rs13479656 | MGI:95522 | Fgfr1 | fibroblast growth factor receptor 1 | protein coding gene | ENSMUSG00000031565 | 14182 | GO:0001525 | angiogenesis |
| rs13479656 | MGI:95522 | Fgfr1 | fibroblast growth factor receptor 1 | protein coding gene | ENSMUSG00000031565 | 14182 | GO:0005524 | ATP binding |
| rs13479656 | MGI:95522 | Fgfr1 | fibroblast growth factor receptor 1 | protein coding gene | ENSMUSG00000031565 | 14182 | GO:0060117 | auditory receptor cell development |
| rs13479656 | MGI:95522 | Fgfr1 | fibroblast growth factor receptor 1 | protein coding gene | ENSMUSG00000031565 | 14182 | GO:0048514 | blood vessel morphogenesis |
| rs13479656 | MGI:95522 | Fgfr1 | fibroblast growth factor receptor 1 | protein coding gene | ENSMUSG00000031565 | 14182 | GO:0007420 | brain development |
| rs13479656 | MGI:95522 | Fgfr1 | fibroblast growth factor receptor 1 | protein coding gene | ENSMUSG00000031565 | 14182 | GO:0060445 | branching involved in salivary gland morphogenesis |
| rs13479656 | MGI:95522 | Fgfr1 | fibroblast growth factor receptor 1 | protein coding gene | ENSMUSG00000031565 | 14182 | GO:0060445 | branching involved in salivary gland morphogenesis |
| rs13479656 | MGI:95522 | Fgfr1 | fibroblast growth factor receptor 1 | protein coding gene | ENSMUSG00000031565 | 14182 | GO:0050839 | cell adhesion molecule binding |
| rs13479656 | MGI:95522 | Fgfr1 | fibroblast growth factor receptor 1 | protein coding gene | ENSMUSG00000031565 | 14182 | GO:0048469 | cell maturation |
| rs13479656 | MGI:95522 | Fgfr1 | fibroblast growth factor receptor 1 | protein coding gene | ENSMUSG00000031565 | 14182 | GO:0021954 | central nervous system neuron development |
| rs13479656 | MGI:95522 | Fgfr1 | fibroblast growth factor receptor 1 | protein coding gene | ENSMUSG00000031565 | 14182 | GO:0002062 | chondrocyte differentiation |
| rs13479656 | MGI:95522 | Fgfr1 | fibroblast growth factor receptor 1 | protein coding gene | ENSMUSG00000031565 | 14182 | GO:0005737 | cytoplasm |
| rs13479656 | MGI:95522 | Fgfr1 | fibroblast growth factor receptor 1 | protein coding gene | ENSMUSG00000031565 | 14182 | GO:0031410 | cytoplasmic vesicle |
| rs13479656 | MGI:95522 | Fgfr1 | fibroblast growth factor receptor 1 | protein coding gene | ENSMUSG00000031565 | 14182 | GO:0043583 | ear development |
| rs13479656 | MGI:95522 | Fgfr1 | fibroblast growth factor receptor 1 | protein coding gene | ENSMUSG00000031565 | 14182 | GO:0030326 | embryonic limb morphogenesis |
| rs13479656 | MGI:95522 | Fgfr1 | fibroblast growth factor receptor 1 | protein coding gene | ENSMUSG00000031565 | 14182 | GO:0001837 | epithelial to mesenchymal transition |
| rs13479656 | MGI:95522 | Fgfr1 | fibroblast growth factor receptor 1 | protein coding gene | ENSMUSG00000031565 | 14182 | GO:0005007 | fibroblast growth factor-activated receptor activity |
| rs13479656 | MGI:95522 | Fgfr1 | fibroblast growth factor receptor 1 | protein coding gene | ENSMUSG00000031565 | 14182 | GO:0005007 | fibroblast growth factor-activated receptor activity |
| rs13479656 | MGI:95522 | Fgfr1 | fibroblast growth factor receptor 1 | protein coding gene | ENSMUSG00000031565 | 14182 | GO:0017134 | fibroblast growth factor binding |
| rs13479656 | MGI:95522 | Fgfr1 | fibroblast growth factor receptor 1 | protein coding gene | ENSMUSG00000031565 | 14182 | GO:0017134 | fibroblast growth factor binding |
| rs13479656 | MGI:95522 | Fgfr1 | fibroblast growth factor receptor 1 | protein coding gene | ENSMUSG00000031565 | 14182 | GO:0017134 | fibroblast growth factor binding |
| rs13479656 | MGI:95522 | Fgfr1 | fibroblast growth factor receptor 1 | protein coding gene | ENSMUSG00000031565 | 14182 | GO:0008543 | fibroblast growth factor receptor signaling pathway |
| rs13479656 | MGI:95522 | Fgfr1 | fibroblast growth factor receptor 1 | protein coding gene | ENSMUSG00000031565 | 14182 | GO:0008543 | fibroblast growth factor receptor signaling pathway |
| rs13479656 | MGI:95522 | Fgfr1 | fibroblast growth factor receptor 1 | protein coding gene | ENSMUSG00000031565 | 14182 | GO:0008543 | fibroblast growth factor receptor signaling pathway |
| rs13479656 | MGI:95522 | Fgfr1 | fibroblast growth factor receptor 1 | protein coding gene | ENSMUSG00000031565 | 14182 | GO:0008543 | fibroblast growth factor receptor signaling pathway |
| rs13479656 | MGI:95522 | Fgfr1 | fibroblast growth factor receptor 1 | protein coding gene | ENSMUSG00000031565 | 14182 | GO:0008543 | fibroblast growth factor receptor signaling pathway |
| rs13479656 | MGI:95522 | Fgfr1 | fibroblast growth factor receptor 1 | protein coding gene | ENSMUSG00000031565 | 14182 | GO:0035607 | fibroblast growth factor receptor signaling pathway involved in orbitofrontal cortex development |
| rs13479656 | MGI:95522 | Fgfr1 | fibroblast growth factor receptor 1 | protein coding gene | ENSMUSG00000031565 | 14182 | GO:0048699 | generation of neurons |
| rs13479656 | MGI:95522 | Fgfr1 | fibroblast growth factor receptor 1 | protein coding gene | ENSMUSG00000031565 | 14182 | GO:0008201 | heparin binding |
| rs13479656 | MGI:95522 | Fgfr1 | fibroblast growth factor receptor 1 | protein coding gene | ENSMUSG00000031565 | 14182 | GO:0042802 | identical protein binding |
| rs13479656 | MGI:95522 | Fgfr1 | fibroblast growth factor receptor 1 | protein coding gene | ENSMUSG00000031565 | 14182 | GO:0042472 | inner ear morphogenesis |
| rs13479656 | MGI:95522 | Fgfr1 | fibroblast growth factor receptor 1 | protein coding gene | ENSMUSG00000031565 | 14182 | GO:0016021 | integral component of membrane |
| rs13479656 | MGI:95522 | Fgfr1 | fibroblast growth factor receptor 1 | protein coding gene | ENSMUSG00000031565 | 14182 | GO:0001701 | in utero embryonic development |
| rs13479656 | MGI:95522 | Fgfr1 | fibroblast growth factor receptor 1 | protein coding gene | ENSMUSG00000031565 | 14182 | GO:0016301 | kinase activity |
| rs13479656 | MGI:95522 | Fgfr1 | fibroblast growth factor receptor 1 | protein coding gene | ENSMUSG00000031565 | 14182 | GO:0060484 | lung-associated mesenchyme development |
| rs13479656 | MGI:95522 | Fgfr1 | fibroblast growth factor receptor 1 | protein coding gene | ENSMUSG00000031565 | 14182 | GO:0030324 | lung development |
| rs13479656 | MGI:95522 | Fgfr1 | fibroblast growth factor receptor 1 | protein coding gene | ENSMUSG00000031565 | 14182 | GO:0030324 | lung development |
| rs13479656 | MGI:95522 | Fgfr1 | fibroblast growth factor receptor 1 | protein coding gene | ENSMUSG00000031565 | 14182 | GO:0016020 | membrane |
| rs13479656 | MGI:95522 | Fgfr1 | fibroblast growth factor receptor 1 | protein coding gene | ENSMUSG00000031565 | 14182 | GO:0048762 | mesenchymal cell differentiation |
| rs13479656 | MGI:95522 | Fgfr1 | fibroblast growth factor receptor 1 | protein coding gene | ENSMUSG00000031565 | 14182 | GO:0030901 | midbrain development |
| rs13479656 | MGI:95522 | Fgfr1 | fibroblast growth factor receptor 1 | protein coding gene | ENSMUSG00000031565 | 14182 | GO:0030901 | midbrain development |
| rs13479656 | MGI:95522 | Fgfr1 | fibroblast growth factor receptor 1 | protein coding gene | ENSMUSG00000031565 | 14182 | GO:0042474 | middle ear morphogenesis |
| rs13479656 | MGI:95522 | Fgfr1 | fibroblast growth factor receptor 1 | protein coding gene | ENSMUSG00000031565 | 14182 | GO:0021837 | motogenic signaling involved in postnatal olfactory bulb interneuron migration |
| rs13479656 | MGI:95522 | Fgfr1 | fibroblast growth factor receptor 1 | protein coding gene | ENSMUSG00000031565 | 14182 | GO:0090272 | negative regulation of fibroblast growth factor production |
| rs13479656 | MGI:95522 | Fgfr1 | fibroblast growth factor receptor 1 | protein coding gene | ENSMUSG00000031565 | 14182 | GO:0010629 | negative regulation of gene expression |
| rs13479656 | MGI:95522 | Fgfr1 | fibroblast growth factor receptor 1 | protein coding gene | ENSMUSG00000031565 | 14182 | GO:0010629 | negative regulation of gene expression |
| rs13479656 | MGI:95522 | Fgfr1 | fibroblast growth factor receptor 1 | protein coding gene | ENSMUSG00000031565 | 14182 | GO:0045668 | negative regulation of osteoblast differentiation |
| rs13479656 | MGI:95522 | Fgfr1 | fibroblast growth factor receptor 1 | protein coding gene | ENSMUSG00000031565 | 14182 | GO:0000122 | negative regulation of transcription by RNA polymerase II |
| rs13479656 | MGI:95522 | Fgfr1 | fibroblast growth factor receptor 1 | protein coding gene | ENSMUSG00000031565 | 14182 | GO:0031175 | neuron projection development |
| rs13479656 | MGI:95522 | Fgfr1 | fibroblast growth factor receptor 1 | protein coding gene | ENSMUSG00000031565 | 14182 | GO:0000166 | nucleotide binding |
| rs13479656 | MGI:95522 | Fgfr1 | fibroblast growth factor receptor 1 | protein coding gene | ENSMUSG00000031565 | 14182 | GO:0005634 | nucleus |
| rs13479656 | MGI:95522 | Fgfr1 | fibroblast growth factor receptor 1 | protein coding gene | ENSMUSG00000031565 | 14182 | GO:0021769 | orbitofrontal cortex development |
| rs13479656 | MGI:95522 | Fgfr1 | fibroblast growth factor receptor 1 | protein coding gene | ENSMUSG00000031565 | 14182 | GO:0001759 | organ induction |
| rs13479656 | MGI:95522 | Fgfr1 | fibroblast growth factor receptor 1 | protein coding gene | ENSMUSG00000031565 | 14182 | GO:0042473 | outer ear morphogenesis |
| rs13479656 | MGI:95522 | Fgfr1 | fibroblast growth factor receptor 1 | protein coding gene | ENSMUSG00000031565 | 14182 | GO:0048339 | paraxial mesoderm development |
| rs13479656 | MGI:95522 | Fgfr1 | fibroblast growth factor receptor 1 | protein coding gene | ENSMUSG00000031565 | 14182 | GO:0018108 | peptidyl-tyrosine phosphorylation |
| rs13479656 | MGI:95522 | Fgfr1 | fibroblast growth factor receptor 1 | protein coding gene | ENSMUSG00000031565 | 14182 | GO:0018108 | peptidyl-tyrosine phosphorylation |
| rs13479656 | MGI:95522 | Fgfr1 | fibroblast growth factor receptor 1 | protein coding gene | ENSMUSG00000031565 | 14182 | GO:0048471 | perinuclear region of cytoplasm |
| rs13479656 | MGI:95522 | Fgfr1 | fibroblast growth factor receptor 1 | protein coding gene | ENSMUSG00000031565 | 14182 | GO:0016310 | phosphorylation |
| rs13479656 | MGI:95522 | Fgfr1 | fibroblast growth factor receptor 1 | protein coding gene | ENSMUSG00000031565 | 14182 | GO:0005886 | plasma membrane |
| rs13479656 | MGI:95522 | Fgfr1 | fibroblast growth factor receptor 1 | protein coding gene | ENSMUSG00000031565 | 14182 | GO:0005886 | plasma membrane |
| rs13479656 | MGI:95522 | Fgfr1 | fibroblast growth factor receptor 1 | protein coding gene | ENSMUSG00000031565 | 14182 | GO:0043536 | positive regulation of blood vessel endothelial cell migration |
| rs13479656 | MGI:95522 | Fgfr1 | fibroblast growth factor receptor 1 | protein coding gene | ENSMUSG00000031565 | 14182 | GO:0060045 | positive regulation of cardiac muscle cell proliferation |
| rs13479656 | MGI:95522 | Fgfr1 | fibroblast growth factor receptor 1 | protein coding gene | ENSMUSG00000031565 | 14182 | GO:0060045 | positive regulation of cardiac muscle cell proliferation |
| rs13479656 | MGI:95522 | Fgfr1 | fibroblast growth factor receptor 1 | protein coding gene | ENSMUSG00000031565 | 14182 | GO:0060045 | positive regulation of cardiac muscle cell proliferation |
| rs13479656 | MGI:95522 | Fgfr1 | fibroblast growth factor receptor 1 | protein coding gene | ENSMUSG00000031565 | 14182 | GO:0045787 | positive regulation of cell cycle |
| rs13479656 | MGI:95522 | Fgfr1 | fibroblast growth factor receptor 1 | protein coding gene | ENSMUSG00000031565 | 14182 | GO:0008284 | positive regulation of cell population proliferation |
| rs13479656 | MGI:95522 | Fgfr1 | fibroblast growth factor receptor 1 | protein coding gene | ENSMUSG00000031565 | 14182 | GO:0008284 | positive regulation of cell population proliferation |
| rs13479656 | MGI:95522 | Fgfr1 | fibroblast growth factor receptor 1 | protein coding gene | ENSMUSG00000031565 | 14182 | GO:0008284 | positive regulation of cell population proliferation |
| rs13479656 | MGI:95522 | Fgfr1 | fibroblast growth factor receptor 1 | protein coding gene | ENSMUSG00000031565 | 14182 | GO:0008284 | positive regulation of cell population proliferation |
| rs13479656 | MGI:95522 | Fgfr1 | fibroblast growth factor receptor 1 | protein coding gene | ENSMUSG00000031565 | 14182 | GO:0008284 | positive regulation of cell population proliferation |
| rs13479656 | MGI:95522 | Fgfr1 | fibroblast growth factor receptor 1 | protein coding gene | ENSMUSG00000031565 | 14182 | GO:0008284 | positive regulation of cell population proliferation |
| rs13479656 | MGI:95522 | Fgfr1 | fibroblast growth factor receptor 1 | protein coding gene | ENSMUSG00000031565 | 14182 | GO:2000546 | positive regulation of endothelial cell chemotaxis to fibroblast growth factor |
| rs13479656 | MGI:95522 | Fgfr1 | fibroblast growth factor receptor 1 | protein coding gene | ENSMUSG00000031565 | 14182 | GO:0010763 | positive regulation of fibroblast migration |
| rs13479656 | MGI:95522 | Fgfr1 | fibroblast growth factor receptor 1 | protein coding gene | ENSMUSG00000031565 | 14182 | GO:2000491 | positive regulation of hepatic stellate cell activation |
| rs13479656 | MGI:95522 | Fgfr1 | fibroblast growth factor receptor 1 | protein coding gene | ENSMUSG00000031565 | 14182 | GO:0050729 | positive regulation of inflammatory response |
| rs13479656 | MGI:95522 | Fgfr1 | fibroblast growth factor receptor 1 | protein coding gene | ENSMUSG00000031565 | 14182 | GO:0043410 | positive regulation of MAPK cascade |
| rs13479656 | MGI:95522 | Fgfr1 | fibroblast growth factor receptor 1 | protein coding gene | ENSMUSG00000031565 | 14182 | GO:0043406 | positive regulation of MAP kinase activity |
| rs13479656 | MGI:95522 | Fgfr1 | fibroblast growth factor receptor 1 | protein coding gene | ENSMUSG00000031565 | 14182 | GO:0090080 | positive regulation of MAPKKK cascade by fibroblast growth factor receptor signaling pathway |
| rs13479656 | MGI:95522 | Fgfr1 | fibroblast growth factor receptor 1 | protein coding gene | ENSMUSG00000031565 | 14182 | GO:0002053 | positive regulation of mesenchymal cell proliferation |
| rs13479656 | MGI:95522 | Fgfr1 | fibroblast growth factor receptor 1 | protein coding gene | ENSMUSG00000031565 | 14182 | GO:0002053 | positive regulation of mesenchymal cell proliferation |
| rs13479656 | MGI:95522 | Fgfr1 | fibroblast growth factor receptor 1 | protein coding gene | ENSMUSG00000031565 | 14182 | GO:0002053 | positive regulation of mesenchymal cell proliferation |
| rs13479656 | MGI:95522 | Fgfr1 | fibroblast growth factor receptor 1 | protein coding gene | ENSMUSG00000031565 | 14182 | GO:1903465 | positive regulation of mitotic cell cycle DNA replication |
| rs13479656 | MGI:95522 | Fgfr1 | fibroblast growth factor receptor 1 | protein coding gene | ENSMUSG00000031565 | 14182 | GO:0045666 | positive regulation of neuron differentiation |
| rs13479656 | MGI:95522 | Fgfr1 | fibroblast growth factor receptor 1 | protein coding gene | ENSMUSG00000031565 | 14182 | GO:0010976 | positive regulation of neuron projection development |
| rs13479656 | MGI:95522 | Fgfr1 | fibroblast growth factor receptor 1 | protein coding gene | ENSMUSG00000031565 | 14182 | GO:0010976 | positive regulation of neuron projection development |
| rs13479656 | MGI:95522 | Fgfr1 | fibroblast growth factor receptor 1 | protein coding gene | ENSMUSG00000031565 | 14182 | GO:2000830 | positive regulation of parathyroid hormone secretion |
| rs13479656 | MGI:95522 | Fgfr1 | fibroblast growth factor receptor 1 | protein coding gene | ENSMUSG00000031565 | 14182 | GO:0010863 | positive regulation of phospholipase C activity |
| rs13479656 | MGI:95522 | Fgfr1 | fibroblast growth factor receptor 1 | protein coding gene | ENSMUSG00000031565 | 14182 | GO:0051897 | positive regulation of protein kinase B signaling |
| rs13479656 | MGI:95522 | Fgfr1 | fibroblast growth factor receptor 1 | protein coding gene | ENSMUSG00000031565 | 14182 | GO:0045944 | positive regulation of transcription by RNA polymerase II |
| rs13479656 | MGI:95522 | Fgfr1 | fibroblast growth factor receptor 1 | protein coding gene | ENSMUSG00000031565 | 14182 | GO:1905564 | positive regulation of vascular endothelial cell proliferation |
| rs13479656 | MGI:95522 | Fgfr1 | fibroblast growth factor receptor 1 | protein coding gene | ENSMUSG00000031565 | 14182 | GO:0046777 | protein autophosphorylation |
| rs13479656 | MGI:95522 | Fgfr1 | fibroblast growth factor receptor 1 | protein coding gene | ENSMUSG00000031565 | 14182 | GO:0005515 | protein binding |
| rs13479656 | MGI:95522 | Fgfr1 | fibroblast growth factor receptor 1 | protein coding gene | ENSMUSG00000031565 | 14182 | GO:0044877 | protein-containing complex binding |
| rs13479656 | MGI:95522 | Fgfr1 | fibroblast growth factor receptor 1 | protein coding gene | ENSMUSG00000031565 | 14182 | GO:0042803 | protein homodimerization activity |
| rs13479656 | MGI:95522 | Fgfr1 | fibroblast growth factor receptor 1 | protein coding gene | ENSMUSG00000031565 | 14182 | GO:0004672 | protein kinase activity |
| rs13479656 | MGI:95522 | Fgfr1 | fibroblast growth factor receptor 1 | protein coding gene | ENSMUSG00000031565 | 14182 | GO:0006468 | protein phosphorylation |
| rs13479656 | MGI:95522 | Fgfr1 | fibroblast growth factor receptor 1 | protein coding gene | ENSMUSG00000031565 | 14182 | GO:0004713 | protein tyrosine kinase activity |
| rs13479656 | MGI:95522 | Fgfr1 | fibroblast growth factor receptor 1 | protein coding gene | ENSMUSG00000031565 | 14182 | GO:0043235 | receptor complex |
| rs13479656 | MGI:95522 | Fgfr1 | fibroblast growth factor receptor 1 | protein coding gene | ENSMUSG00000031565 | 14182 | GO:0090722 | receptor-receptor interaction |
| rs13479656 | MGI:95522 | Fgfr1 | fibroblast growth factor receptor 1 | protein coding gene | ENSMUSG00000031565 | 14182 | GO:0060665 | regulation of branching involved in salivary gland morphogenesis by mesenchymal-epithelial signaling |
| rs13479656 | MGI:95522 | Fgfr1 | fibroblast growth factor receptor 1 | protein coding gene | ENSMUSG00000031565 | 14182 | GO:0042127 | regulation of cell population proliferation |
| rs13479656 | MGI:95522 | Fgfr1 | fibroblast growth factor receptor 1 | protein coding gene | ENSMUSG00000031565 | 14182 | GO:0042127 | regulation of cell population proliferation |
| rs13479656 | MGI:95522 | Fgfr1 | fibroblast growth factor receptor 1 | protein coding gene | ENSMUSG00000031565 | 14182 | GO:2001239 | regulation of extrinsic apoptotic signaling pathway in absence of ligand |
| rs13479656 | MGI:95522 | Fgfr1 | fibroblast growth factor receptor 1 | protein coding gene | ENSMUSG00000031565 | 14182 | GO:0010468 | regulation of gene expression |
| rs13479656 | MGI:95522 | Fgfr1 | fibroblast growth factor receptor 1 | protein coding gene | ENSMUSG00000031565 | 14182 | GO:0048378 | regulation of lateral mesodermal cell fate specification |
| rs13479656 | MGI:95522 | Fgfr1 | fibroblast growth factor receptor 1 | protein coding gene | ENSMUSG00000031565 | 14182 | GO:0010966 | regulation of phosphate transport |
| rs13479656 | MGI:95522 | Fgfr1 | fibroblast growth factor receptor 1 | protein coding gene | ENSMUSG00000031565 | 14182 | GO:0051174 | regulation of phosphorus metabolic process |
| rs13479656 | MGI:95522 | Fgfr1 | fibroblast growth factor receptor 1 | protein coding gene | ENSMUSG00000031565 | 14182 | GO:0051930 | regulation of sensory perception of pain |
| rs13479656 | MGI:95522 | Fgfr1 | fibroblast growth factor receptor 1 | protein coding gene | ENSMUSG00000031565 | 14182 | GO:0072091 | regulation of stem cell proliferation |
| rs13479656 | MGI:95522 | Fgfr1 | fibroblast growth factor receptor 1 | protein coding gene | ENSMUSG00000031565 | 14182 | GO:0007435 | salivary gland morphogenesis |
| rs13479656 | MGI:95522 | Fgfr1 | fibroblast growth factor receptor 1 | protein coding gene | ENSMUSG00000031565 | 14182 | GO:0007605 | sensory perception of sound |
| rs13479656 | MGI:95522 | Fgfr1 | fibroblast growth factor receptor 1 | protein coding gene | ENSMUSG00000031565 | 14182 | GO:0042169 | SH2 domain binding |
| rs13479656 | MGI:95522 | Fgfr1 | fibroblast growth factor receptor 1 | protein coding gene | ENSMUSG00000031565 | 14182 | GO:0005102 | signaling receptor binding |
| rs13479656 | MGI:95522 | Fgfr1 | fibroblast growth factor receptor 1 | protein coding gene | ENSMUSG00000031565 | 14182 | GO:0019827 | stem cell population maintenance |
| rs13479656 | MGI:95522 | Fgfr1 | fibroblast growth factor receptor 1 | protein coding gene | ENSMUSG00000031565 | 14182 | GO:0016740 | transferase activity |
| rs13479656 | MGI:95522 | Fgfr1 | fibroblast growth factor receptor 1 | protein coding gene | ENSMUSG00000031565 | 14182 | GO:0001657 | ureteric bud development |
| rs13479656 | MGI:95522 | Fgfr1 | fibroblast growth factor receptor 1 | protein coding gene | ENSMUSG00000031565 | 14182 | GO:0060979 | vasculogenesis involved in coronary vascular morphogenesis |
| rs13479656 | MGI:95522 | Fgfr1 | fibroblast growth factor receptor 1 | protein coding gene | ENSMUSG00000031565 | 14182 | GO:0021847 | ventricular zone neuroblast division |
| rs13479656 | MGI:95522 | Fgfr1 | fibroblast growth factor receptor 1 | protein coding gene | ENSMUSG00000031565 | 14182 | GO:0070640 | vitamin D3 metabolic process |
| rs13479656 | MGI:5590447 | Gm31288 | predicted gene, 31288 | lncRNA gene | | 102633471 |  |  |
| rs13479656 | MGI:6004449 | Tssr76682 | transcription start site region 76682 | TSS region |  |  |  |  |
| rs3704385 | MGI:5590943 | Gm31784 | predicted gene, 31784 | lncRNA gene | ENSMUSG00000110229 | 102634123 |  |  |
| rs3704385 | MGI:5791143 | Gm45307 | predicted gene 45307 | lncRNA gene | ENSMUSG00000109682 | |  |  |
| rs13479753 | MGI:1922799 | 1700019L22Rik | RIKEN cDNA 1700019L22 gene | lncRNA gene | | 75549 |  |  |
| rs3664869 | No associated gene | |  |  |  |  |  |  |
| rs3663506 | No associated gene | |  |  |  |  |  |  |
| rs3721390 | No associated gene | |  |  |  |  |  |  |
| rs13479993 | No associated gene | |  |  |  |  |  |  |
| rs13480205 | No associated gene | |  |  |  |  |  |  |
| rs3670195 | MGI:6014103 | Tssr86336 | transcription start site region 86336 | TSS region |  |  |  |  |
